# Structural and mechanistic basis of σ-dependent transcriptional pausing

**DOI:** 10.1101/2022.01.24.477500

**Authors:** Chirangini Pukhrambam, Vadim Molodtsov, Mahdi Kooshbaghi, Ammar Tareen, Hoa Vu, Kyle S. Skalenko, Min Su, Yin Zhou, Jared T. Winkelman, Justin B. Kinney, Richard H. Ebright, Bryce E. Nickels

## Abstract

In σ-dependent transcriptional pausing, the transcription initiation factor σ, translocating with RNA polymerase (RNAP), makes sequence-specific protein-DNA interactions with a promoter-like sequence element in the transcribed region, inducing pausing. It has been proposed that, in σ-dependent pausing, the RNAP active center can access off-pathway “backtracked” states that are substrates for the transcript-cleavage factors of the Gre family, and on-pathway “scrunched” states that mediate pause escape. Here, using site-specific protein-DNA photocrosslinking to define positions of the RNAP trailing and leading edges and of σ relative to DNA at the *λ*PR’ promoter, we show directly that σ-dependent pausing in the absence of GreB *in vitro* predominantly involves a state backtracked by 2-4 bp, and that σ-dependent pausing in the presence of GreB *in vitro* and *in vivo* predominantly involves a state scrunched by 2-3 bp. Analogous experiments with a library of 4^7^ (∼16,000) transcribed-region sequences show that the state scrunched by 2-3 bp--and only that state--is associated with the consensus sequence, T_-3_N_-2_Y_-1_G_+1_, (where -1 corresponds to the position of the RNA 3’ end), which is identical to the consensus for pausing in initial transcription, and which is related to the consensus for pausing in transcription elongation. Experiments with heteroduplex templates show that sequence information at position T_-3_ resides in the DNA nontemplate strand. A cryo-EM structure of a complex engaged in σ-dependent pausing reveals positions of DNA scrunching on the DNA nontemplate and template strands and suggests that position T_-3_ of the consensus sequence exerts its effects by facilitating scrunching.

## Introduction

The RNA polymerase (RNAP) holoenzyme initiates transcription by binding double-stranded promoter DNA, unwinding a turn of promoter DNA to yield an RNAP-promoter open complex (RPo) containing a ∼13 base pair unwound “transcription bubble” and selecting a transcription start site (1–4). In bacteria, promoter binding and promoter unwinding are mediated by the transcription initiation factor σ, which, in the context of the RNAP holoenzyme, participates in sequence-specific protein-DNA interactions with the promoter -35 element, recognized by σ region 4 (σR4), and the promoter -10 element, recognized by σ region 2 (σR2) (5, 6).

The first ∼11 nucleotides (nt) of an RNA product are synthesized as an RNAP-promoter initial transcribing complex (ITC) in which RNAP remains anchored on promoter DNA through sequence-specific interactions with σ (1, 3). In initial transcription, RNAP uses a “scrunching” mechanism of RNAP-active-center translocation, in which, in each nucleotide-addition cycle, RNAP remains stationary on DNA and unwinds one base pair of DNA downstream of the RNAP active center, pulls the unwound single-stranded DNA (ssDNA) into and past the RNAP active center, and accommodates the additional unwound ssDNA as bulges in the transcription bubble (1, 7–10). Scrunching enables the capture of free energy from multiple nucleotide additions and the stepwise storage of captured free energy in the form of stepwise increases in the amount of DNA unwinding (RPitc,2 to RPitc,11). Thus, scrunching provides the mechanism to capture and store the free energy required to break RNAP-promoter interactions in subsequent promoter escape (1, 7, 9, 10).

Following synthesis of an RNA product of ∼11 nt, promoter escape occurs. Promoter escape entails: (i) entry of the RNA 5ʹ end into the RNAP RNA-exit channel; (ii) displacement of σ from the RNAP RNA-exit channel, driven by steric clash with the RNA 5ʹ end (11–14); (iii) disruption of protein-DNA interaction between σ and the promoter -35 element; and (iv) rewinding of the upstream half of the transcription bubble, from the -10 element through the transcription start site, a process termed “unscrunching” (9, 14, 15). The product of this series of reactions is a transcription elongation complex (TEC) containing a threshold length of ∼11 nt of RNA and having an altered RNAP-σ interface in which a subset of the interactions previously made between RNAP and σ are lost (16–19). Because of the partial loss of RNAP σ interactions, the affinity of RNAP for σ is decreased and σ typically dissociates in a time dependent fashion (16, 17, 20–28).

In contrast to initial transcription, which proceeds through a scrunching mechanism, transcription elongation proceeds through a “stepping” mechanism, in which RNAP steps forward by 1 bp relative to DNA for each nucleotide added to the RNA product (29). Each nucleotide-addition cycle of transcription requires translocation of the RNAP active center relative to DNA and RNA, starting from a “pre-translocated” state, and yielding a “post-translocated” state. Translocation of the RNAP active center repositions the RNA 3ʹ end from the RNAP addition site (A site) to the RNAP product site (P site), rendering the A site available to bind the next extending NTP (30–33).

The rate of RNA synthesis is not uniform across the DNA template. At certain template positions, transcription is interrupted by “pausing”-- i.e., nucleotide-addition cycles that occur on the second or longer timescale (3, 30, 32, 34). Pausing can often involve entry of the transcription complex into an off pathway, backtracked state where the RNAP active center has reverse translocated relative to DNA and RNA, rendering the active center unable to add NTPs (35–38). Pausing can impact gene expression by reducing the rate of RNA synthesis, facilitating engagement of regulatory factors with RNAP, modulating formation of RNA secondary structures, or enabling synchronization of transcription and translation (39). RNAP can be induced to pause by DNA sequences (sequence-dependent pausing) or by interacting proteins (factor-dependent pausing)(34).

The first identified and still paradigmatic example of factor-dependent pausing is σ-dependent pausing (16, 40–43). A σ-containing TEC in a promoter-proximal transcribed region (prior to σ dissociation), or, to a lesser degree, in a promoter-distal transcribed region (following σ dissociation and σ reassociation), can recognize and engage, through sequence-specific σ-DNA interactions, a transcribed-region sequence that resembles a promoter element (16, 21, 22, 41–48). These sequence-specific σ-DNA interactions anchor the σ-containing TEC at the sequence resembling a promoter element, resulting in σ-dependent transcriptional pausing. Typically, a σ-dependent pause element (SDPE) is a sequence that resembles a consensus promoter -10 element, often supplemented by a sequence that resembles a consensus discriminator element. σ-dependent pausing, particularly σ-dependent pausing in promoter-proximal regions, enables coordination of the timing of transcription elongation with the timing of other biological processes. In the best-characterized example, σ-dependent pausing 16-17 bp downstream of the transcription start site of the bacteriophage *λ* PR’ promoter (*λ*PR’; Figure 1A) coordinates transcription elongation and regulation of transcription termination, by providing time for loading of the transcription antitermination factor *λ*Q (40, 49–51). In other well-characterized examples, σ-dependent pausing 18 bp and 25 bp downstream of the bacteriophage 21 and 82 PR’ promoters coordinates transcription elongation and regulation of transcription termination in a similar manner (41, 49, 52, 53). Genome-wide analyses suggest that σ-dependent pausing occurs in as many as 20% of transcription units in *E. coli* (45, 54) and is functionally linked to expression levels of stress-related genes (54).

**Figure 1.**
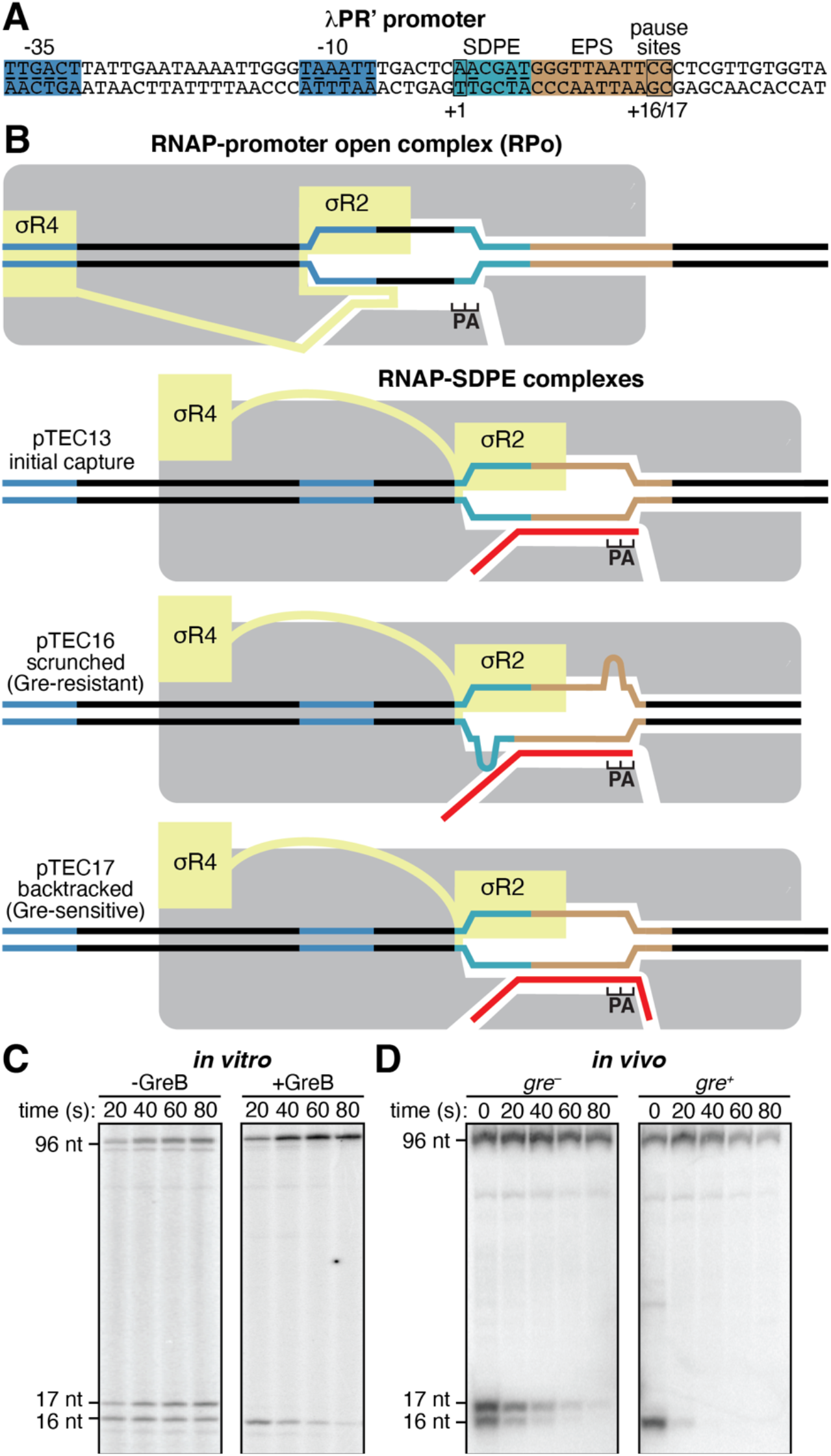
σ-dependent pausing at λPR’: scrunched and backtracked states. (A) *λ*PR’ promoter. Blue, -35-element and -10-element; light blue, SDPE; brown; elemental pause site (EPS); black rectangles, transcription start site (+1) and pause sites (+16/+17); underlining, consensus nucleotides of sequence elements. (B) Initiation complexes and paused complexes at *λ*PR’. Four complexes are shown: (1) initiation complex, RPo; (2) initial-capture σ-dependent paused complex, pTEC13 (where “pTEC” denotes paused TEC and “13” denotes 13 nt RNA product); (3) scrunched σ-dependent paused complex, pTEC16; and, (4) backtracked σ-dependent paused complex, pTEC17. Gray, RNAP core; yellow, *σ*; red, RNA product; P and A, RNAP active-center product and addition sites; blue, -35-element and -10-element; light blue, SDPE; brown, EPS; black, other DNA (nontemplate-strand above template-strand). Scrunching of DNA strands is indicated by bulges in DNA strands. (C-D) RNA product length and Gre-factor sensitivity in pTECs. Panel C shows RNA product distributions *in vitro* for transcription reactions in absence or presence of GreB at indicated times after addition of NTPs. Panel D shows RNA product distributions *in vivo* for *gre^-^* or *gre^+^* cells at indicated times after addition of rifampin (Rif). Positions of pTEC-associated RNA products (16 nt and 17 nt) and full-length RNA product (96 nt) are indicated.

It has been proposed that in σ-dependent pausing, following the initial engagement of the SDPE (“pause capture”; Figure 1B, line 2), the paused TEC (pTEC) can extend RNA for several nucleotides, using a scrunching mechanism (Figure 1B, line 3), and that the resulting pTECs with extended RNA equilibrate between scrunched states and backtracked states (Figure 1B, lines 3-4) (10, 41, 55–59). It further has been proposed that the scrunched states are intermediates on the pathway to pause escape, that the backtracked states are off the pathway to pause escape, and that DNA sequences downstream of the SDPE, including sequences related to the consensus sequence for elemental pausing in transcription elongation, modulate the duration of the pause (“pause lifetime”) through effects on the relative occupancies of scrunched and backtracked states (41, 55–59). Evidence in support of these proposals comes from measurement of RNA product lengths, DNA footprinting (40, 41, 50, 55–58), and analysis of sensitivity of complexes to transcript cleavage factors of the Gre family (45–47, 55) (which promote transcript cleavage in backtracked states but not in other states; Figure 1C; (60)). However, the evidence is not dispositive.

Here, we use *in vitro* and *in vivo* site-specific protein-DNA photocrosslinking, high-throughput sequencing, heteroduplex-template transcription experiments, and cryo-EM structure determination, to define the mechanistic and structural basis of σ-dependent pausing at *λ*PR’.

## Results

### σ-dependent pausing at λ PR’: pTEC RNA-product length and Gre-factor sensitivity

It previously has been shown that at *λ*PR’, *in vitro*, in the absence of Gre factors, the pTEC is present both in a state with 16 nt of RNA (pTEC16) and a state with 17 nt of RNA (pTEC17) (40, 51, 55). It also previously has been shown that at *λ*PR’, *in vitro*, in the presence of Gre factors, the pTEC is present predominantly as pTEC16 (55). The data in Figure 1C confirm these results and the data in Figure 1D show that the same pattern is obtained *in vivo*. Thus, in *gre*^-^ cells, the pTEC is present both as pTEC16 and as pTEC17, whereas in *gre*^+^ cells, the pTEC is present essentially exclusively as pTEC16.

### Use of site-specific protein-DNA photocrosslinking to define positions of RNAP trailing and leading edges and of σ relative to DNA at λPR’: approach

We used unnatural amino acid mutagenesis (61) to incorporate the photoactivatable crosslinking agent *p*-benzoyl-L-phenylalanine (Bpa) into RNAP holoenzyme at specific positions on the RNAP trailing edge, the RNAP leading edge, and σR2 (*β*’ residue 48, *β*’ residue 1148, and σ^70^ residue 448; Figure 2). The resulting RNAP holoenzyme derivatives behaved indistinguishably from unmodified wild-type RNAP in terms of RNA-product-length distributions of paused complexes and Gre sensitivities of paused complexes (Figure S1). Using the resulting RNAP holoenzyme derivatives, in purified form for experiments *in vitro*, and *in situ*, inside living cells, for experiments *in vivo*, we performed site-specific protein-DNA photocrosslinking (62–65), to define the positions of the RNAP trailing edge, the RNAP leading edge, and σR2 relative to DNA in RNAP-promoter complexes (RPo) and in RNAP-SDPE complexes (pTEC) at *λ*PR’ (Figures 3 and S2-S4). From the observed positions of the RNAP leading edge relative to DNA, we then inferred the position of the RNAP-active-center nucleotide-addition site (A site) by subtracting 5 nt, as described previously (65).

**Figure 2.**
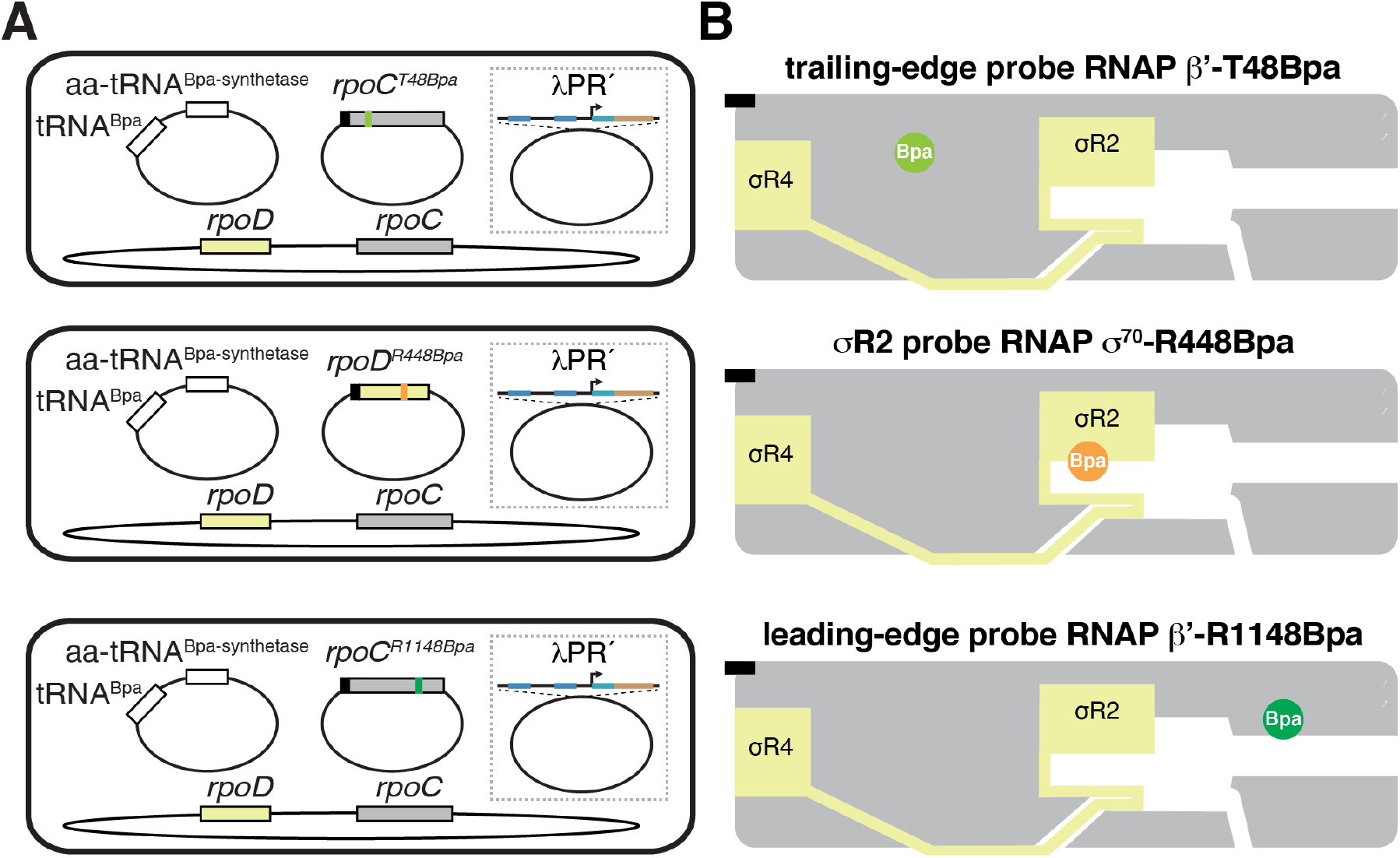
Use of site-specific protein-DNA photocrosslinking to define positions of RNAP trailing and leading edges and of σ relative to DNA at λPR’: approach. (A) Two-plasmid (for *in vitro* studies) or three-plasmid (for *in vivo* studies) merodiploid system for co-production, in *E. coli* cells, of decahistidine-tagged RNAP-*β*’ ^T48Bpa^, RNAP-*β*’ ^R1148Bpa^, or RNAP-*σ*^70 R448Bpa^ in the presence of untagged wild-type RNAP holoenzyme. First plasmid carries gene for decahistidine-tagged RNAP βʹ subunit (gray rectangle) with nonsense codon at position 48 (olive green; top row); decahistidine-tagged *σ*^70^ (light yellow rectangle) with nonsense codon at position 448 (orange; center row); or gene for decahistidine-tagged RNAP βʹ subunit with nonsense codon at position 1148 (forest green; bottom row). Second plasmid carries genes for engineered Bpa-specific nonsense-suppressor tRNA and aminoacyl-tRNA synthetase (white rectangles). Third plasmid (shown inside dashed box), when present, carries *λ*PR’ promoter or *λ*PR’ promoter derivative. Chromosome (shown below plasmids) carries genes for wild-type RNAP βʹ subunit and *σ*^70^. Black rectangles, decahistidine-tag coding sequence. (B) Bpa-modified RNAPs. Olive green circle, trailing-edge Bpa; orange circle, *σ*R2 Bpa; forest green circle, leading-edge Bpa. Black rectangles, decahistidine-tag. Other colors and symbols as in Figure 1A-B.

**Figure 3.**
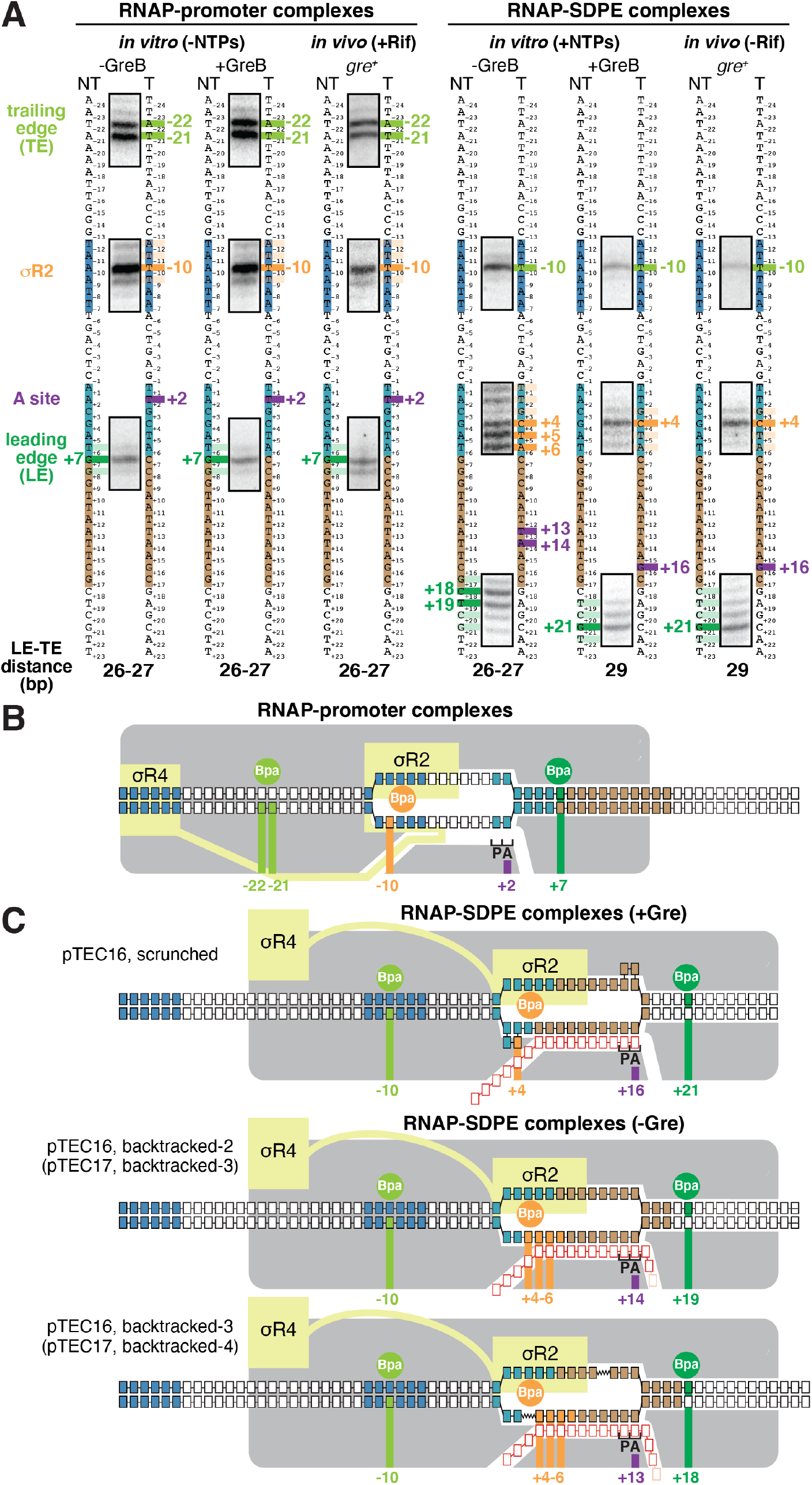
Use of site-specific protein-DNA photocrosslinking to define positions of RNAP trailing and leading edges and of σ relative to DNA at λPR’: results. (A) Positions of RNAP trailing and leading edges and of σR2 in RNAP-promoter complexes (left) or RNAP-SDPE complexes (pTECs) (right) at *λ*PR’. For each experimental condition *in vitro* and *in vivo*, identified at top, figure shows segments of gel images for primer-extension mapping of crosslinking sites (full gel images in Figures S2-S4), nontemplate- and template-strand sequences of *λ*PR’ (to left and right, respectively, of gel images; -35 element, -10 element, SDPE, and EPS colored as in Figure 1A), observed crosslinking sites (olive green for RNAP trailing edge, forest green for RNAP leading edge, and orange for σR2), inferred positions of RNAP- active-center A site (violet), and inferred modal trailing-edge/leading-edge distances (TE-LE distances). (B) Mechanistic interpretation of data for RNAP-promoter complexes at *λ*PR’ (panel A, left). Olive green circle and olive green vertical lines denote Bpa site at RNAP trailing edge and observed crosslinking sites in DNA for Bpa at RNAP trailing edge, forest green circle and forest green vertical line denote Bpa site at RNAP leading edge and observed crosslinking site in DNA for Bpa at RNAP leading edge, and orange circle and orange vertical line denote Bpa site in *σ*R2 and observed crosslinking site in DNA for Bpa in *σ*R2. Violet vertical line denotes inferred position of RNAP-active-center A site. Gray, RNAP core; yellow, *σ*; P and A, RNAP active-center product and addition sites; black boxes with blue fill, -35-element and -10-element nucleotides; black boxes with light blue fill, SDPE nucleotides; black boxes with brown fill, EPS nucleotides; other black boxes, other DNA nucleotides (nontemplate-strand nucleotides above template-strand nucleotides). (C) Mechanistic interpretation of data for RNAP-SDPE complexes at *λ*PR’ (pTEC; panel A, right). Three complexes are shown: (1) a scrunched σ-dependent paused complex with 16 nt RNA product and RNAP-active-center A-site at position +16 (pTEC16 scrunched; top row) (2) a backtracked σ-dependent paused complex with 16 or 17 nt RNA product and RNAP-active-center at position +14 (pTEC16, backtracked-2 or pTEC17, backtracked-3; center row), and (3) a backtracked σ-dependent paused complex with 16 or 17 nt RNA product and RNAP-active-center at position +13 (pTEC16, backtracked-3 or pTEC17, backtracked-4; bottom row). Red boxes, RNA nucleotides; pink boxes, additional RNA nucleotide present in 17 nt RNA product. Other colors as in panel B. Scrunched segments of nontemplate and template DNA strands in pTEC16 shown as bulges.

To assess RPo and pTEC at *λ*PR’ *in vitro*, we formed transcription complexes in the absence and presence, respectively, of nucleoside triphosphates (NTPs) (Figures 3A and S2-S4). To assess RPo and pTEC at *λ*PR’ *in vivo*, we performed experiments in the presence and absence, respectively, of the transcription inhibitor rifampin, which blocks extension of RNA beyond a length of 2-3 nt (66, 67), and thus prevents formation of TEC (Figures 3A and S2-S4).

### Use of site-specific protein-DNA photocrosslinking to define positions of RNAP trailing and leading edges and of σ relative to DNA at λPR’: results

The crosslinking results for RPo at *λ*PR’ show that the RNAP-leading-edge/RNAP-trailing-edge distance (LE-TE distance) is 26-27 bp, consistent with an unscrunched transcription complex (62–64, 68), show that the RNAP active-center A site interacts with promoter position +2, and show that σR2 interacts with the promoter -10 element (promoter position -10) (Figures 3A, left, and 3B). The crosslinking pattern for RPo at *λ*PR’ is identical for experiments performed *in vitro* in the absence of GreB, *in vitro* in the presence of GreB and *in vivo* in *gre*^+^ cells (Figures 3A, left, and 3B).

The crosslinking results for pTEC at *λ*PR’ *in vitro* in the presence of GreB and *in vivo* in *gre*^+^ cells--where, as shown above, the pTEC is present exclusively as pTEC16 (Figure 1C,D)--show that the LE-TE distance is 29 bp, show that the RNAP active-center A site interacts with position +16, and show that σR2 interacts with the SDPE (position +4) (Figure 3A, right, columns 2 and 3). The LE-TE distance in pTEC16 in the presence of Gre indicates that the complex contains 2-3 bp of DNA scrunching (LE-TE distance of 29 bp for pTEC16 vs LE-TE distance of 26-27 bp for RPo). The crosslinking results for the RNAP leading edge in pTEC16 in the presence of Gre indicate that the RNAP active-center A site has translocated forward by 14 steps, from position +2 in RPo to position +16. The crosslinking results for σR2 in pTEC16 in the presence of Gre indicate that σR2 has disengaged from the promoter -10 element and has engaged with the SDPE. Taken together, the crosslinking results establish that pTEC16 in the presence of Gre is a σ-dependent paused TEC having 2-3 bp of DNA scrunching (Figure 3C, top).

The crosslinking results for pTEC at *λ*PR’ *in vitro* in the absence of Gre--where, as shown above, the pTEC is present as both pTEC16 and pTEC17 (Figure 1C)--show that the LE-TE distance is 26-27 bp, show that the RNAP active-center A site interacts with position +13 or +14, and show that σR2 interacts with the SDPE (positions +4 to +6) (Figure 3A, right, column 1). The LE-TE distances in pTEC16 and pTEC17 in the absence of Gre indicate that the complexes are unscrunched (LE-TE distance of 26-27 bp for pTEC16 and pTEC17 in the absence of Gre vs LE-TE distance of 26-27 bp for RPo). The crosslinking results for the RNAP leading edge in pTEC16 and pTEC17 in the absence of Gre together with the 16 and 17 nt lengths of the RNA products in those complexes indicate that the RNAP active-center A site first translocated forward by 14 or 15 steps, from position +2 in RPo to position +16 or +17, and then reverse translocated--backtracked--by 2 to 4 bp. The crosslinking results for σR2 in pTEC16 and pTEC17 in the absence of Gre indicate that σR2 has disengaged from the promoter -10 element and has engaged with the SDPE. Taken together, the crosslinking results establish that pTEC16 and pTEC17 in the absence of Gre are σ-dependent paused TECs backtracked by 2-4 bp (Figure 3C, center and bottom).

The RNAP trailing-edge position in pTEC16 in the presence of Gre is the same as the RNAP trailing-edge positions in pTEC16 and pTEC17 in the absence of Gre (Figure 3A, right, columns 2-3 vs. column 1). In contrast, the RNAP leading-edge position in pTEC16 in the presence of Gre differs by 2-3 bp, in a downstream direction, from the RNAP trailing-edge positions in pTEC16 and pTEC17 in the absence of Gre (Figure 3A, right, columns 2-3 vs. column 1).

A defining hallmark of DNA scrunching is a change in the position of RNAP leading edge relative to DNA without a corresponding change in the position of the RNAP trailing edge relative to DNA, resulting in a change in LE-TE distance (7, 8, 63, 68). The results in Figure 3 establish that σ-dependent pTECs formed *in vitro* or *in vivo* in the presence of Gre exhibit this defining hallmark of DNA scrunching. Thus, the RNAP leading-edge position in pTEC16 in the presence of Gre is different, by 2-3 bp, from the RNAP leading-edge position in pTEC16 in the absence of Gre (Figure 3A, right, columns 2-3 vs. column 1), whereas, in contrast, the RNAP trailing-edge position in pTEC16 in the presence of Gre is identical to the RNAP trailing-edge position in pTEC16 in the absence of Gre (Figure 3A, right, columns 2-3 vs. column 1).

### Sequence determinants for scrunching in σ-dependent pausing

It has been hypothesized that scrunching occurs in σ-dependent pausing and has been further hypothesized that DNA sequences downstream of the SDPE modulate pause lifetime through effects on DNA scrunching (10, 55–59). The results in Figure 3 establish that scrunching occurs in σ-dependent pausing at *λ*PR’ and provide an experimental approach--namely, crosslinking of the RNAP leading edge to DNA--that enables scrunched states and unscrunched states in σ-dependent pausing to be distinguished for any DNA sequence containing an SDPE.

In a next set of experiments, we applied this experimental approach to a library of DNA sequences containing the *λ*PR’ SDPE and all ∼16,000 possible 7 bp DNA sequences spanning the *λ*PR’ pause site (*λ*PR’ positions +14 to +20; Figure 4A) to assess whether DNA sequences downstream of the SDPE determine scrunching in σ-dependent pausing and, if so, to define the sequence determinants (Figure 4).

**Figure 4.**
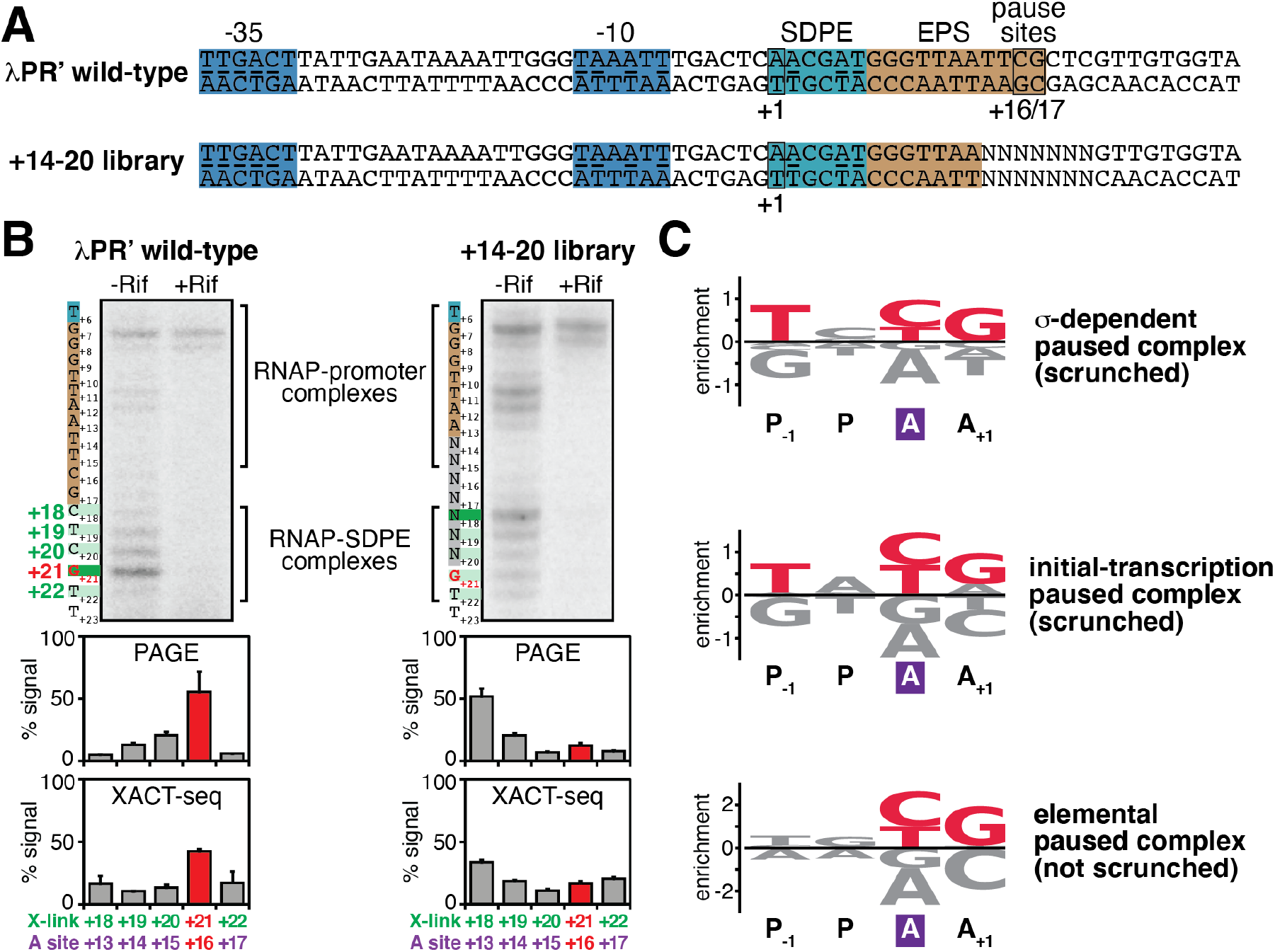
Sequence determinants for scrunching in σ-dependent pausing. (A) DNA templates containing wild-type *λ*PR’ or +14-20 library. NNNNNNN, randomized nucleotides of +14-20 library. Other colors as in Figure 1A. (B) Positions of RNAP leading edge in wild-type *λ*PR’ and +14-20 library *in vivo*. Top subpanel, PAGE analysis of crosslinking. For each experimental condition, identified at top, figure shows gel image for primer-extension mapping of crosslinking sites, nontemplate-strand sequence (to left of gel image; SDPE and EPS colored as in Figure 1A), and observed crosslinking sites (forest green). Center subpanel, quantitation (mean ± SD) of PAGE analysis of crosslinking. Bottom subpanel, quantitation (mean ± SD) of XACT-seq analysis of crosslinking. In all subpanels, the observed major crosslinking site for pTEC at *λ*PR’ (position +21) and inferred major RNAP-active-center A-site position for pTEC at *λ*PR’ (position +16) are highlighted in red. (C) Sequence logos quantifying formation and/or stability of scrunched σ-dependent paused complex (top; this work); formation and/or stability of scrunched initial-transcription paused complex (center; (65)); and elemental pausing in transcription elongation (bottom; (69–72)). Positions are labeled relative to RNAP-active-center A site (violet rectangle) and P-site. Red, most highly preferred DNA nucleotides. Logos were generated using Logomaker (85) as described in SI Materials and Methods.

First, using procedures analogous to those of the previous section, we transformed *gre*^+^ cells producing RNAP holoenzyme having Bpa incorporated at the RNAP leading edge (*β*’ residue 1148) with a plasmid carrying *λ*PR’ or, in parallel, plasmids carrying sequences from the “+14-20 library,” and we then UV-irradiated cells to initiate RNAP-DNA photocrosslinking, lysed cells, isolated crosslinked material, and mapped crosslinks by primer extension and urea-PAGE (Figure 4B). For *λ*PR’, the results show that the RNAP leading-edge crosslinks predominantly at position +21, indicating that the RNAP active-center A site interacts predominantly with positions +16, as expected, based on the results in the preceding section, for the scrunched σ-dependent paused complex at *λ*PR’ (Figures 3 and 4B, left). In contrast, for the +14-20 library, the RNAP leading edge crosslinks predominantly at position +18, indicating that the RNAP active-center A site interacts predominantly with positions +13, as expected for unscrunched complexes (such as, for example, the backtracked σ-dependent paused complexes of Figure 3C). We conclude that positions +14 to +20 of *λ*PR’ contains sequence information crucial for formation and/or stability of the scrunched state during σ-dependent pausing, consistent with previous proposals (57, 58). We further conclude, from comparison of the yield of the scrunched state for the +14-20 library vs. for *λ*PR’ (∼10% vs. ∼50%; Figure 4B, center), that only a fraction, ∼1/5, of the ∼16,000 possible sequences at positions +14 to +20 support formation and/or stability of the scrunched state during σ-dependent pausing.

Second, in order to identify sequence determinants that influence formation and/or stability of the scrunched state during σ-dependent pausing, we performed deep sequencing of the primer-extension products of the preceding paragraph, following the “XACT-seq” procedure of (65) (<u>x</u>linking-of-<u>a</u>ctive-<u>c</u>enter-to-template-<u>seq</u>uencing; Figure 4B, bottom). Consistent with the PAGE analysis of the preceding paragraph, the XACT-seq analysis show that ∼10% of sequence reads corresponded to the scrunched σ-dependent paused complex (i.e., sequence reads for which RNAP-leading-edge crosslinking occurred at position +21, indicating interaction of the RNAP-active-center A site with position +16) (Figure 4B, bottom). Analysis of this subset of sequence reads showed clear sequence preferences, yielding the consensus sequence T_+14_N_+15_Y_+16_G_+17_, or, expressed in terms of RNAP-active-center A-site and P-site positions, T_P-1_N_P_Y_A_G_A+1_ (where N is any nucleotide and Y is pyrimidine) (Figures 4C, top, and S5).

*λ*PR’ contains a perfect match to the consensus sequence T_+14_N_+15_Y_+16_G_+17_ (Figure 4A). *λ*PR’ positions +16 and +17 have been shown to affect pause capture efficiency and pause lifetime (58). Our results indicate that the sequence at positions +16 and +17 affects formation and/or stability of the scrunched state of the σ-dependent paused complex at *λ*PR’, and are consistent with the view that the effects of sequence on pause capture efficiency and pause lifetime are consequences of the effects of sequence on formation and/or stability of the scrunched state.

This consensus sequence obtained in this work for formation and/or stability of a scrunched σ-dependent paused complex (T_P-1_N_P_Y_A_G_A+1_; Figure 4C, top) is identical to the consensus sequence obtained in previous work for formation and/or stability of a scrunched initial-transcription paused complex (T_P-1_N_P_Y_A_G_A+1_; Figure 4C, middle) (65). The consensus sequence also is similar to the downstream, most-highly-conserved, portion of the consensus sequence obtained in previous work for elemental pausing in transcription elongation (Y_A_G_A+1_;

Figure 4C, bottom) (69–72); however, the consensus sequences for the scrunched σ-dependent paused complex and scrunched initial-transcription paused complex show a substantially stronger conservation of T_P-1_ than the consensus sequence for elemental pausing in transcription elongation (Figure 4C).

### Strand-dependence of sequence determinants for scrunching in σ-dependent pausing

As described in the preceding section, the consensus sequence for formation of a scrunched σ-dependent paused complex contains a strongly conserved T:A base pair at position P-1 (T on the nontemplate stand; A on the template strand) that also is strongly conserved in the consensus sequence for formation of a scrunched initial-transcription complex but that is not strongly conserved in the consensus sequence for elemental pausing in transcription elongation (Figure 4C). To determine whether specificity at this position resides in the nontemplate-strand T or the template-strand A, we analyzed formation of the scrunched state in σ-dependent pausing, *in vitro*, in the presence of GreB, using heteroduplex templates in which the nontemplate-strand T was replaced by a non-consensus nucleotide or an abasic site, and the template-strand A was unchanged (Figure 5A). The results show that the presence of a non-consensus nucleotide or an abasic site on the nontemplate strand at position P-1 abrogates formation of the scrunched state in σ-dependent pausing (Figure 5B). We conclude that the sequence information responsible for the preference for T:A at position P-1 resides, at least in part, in the DNA nontemplate strand (Figure 5C).

**Figure 5.**
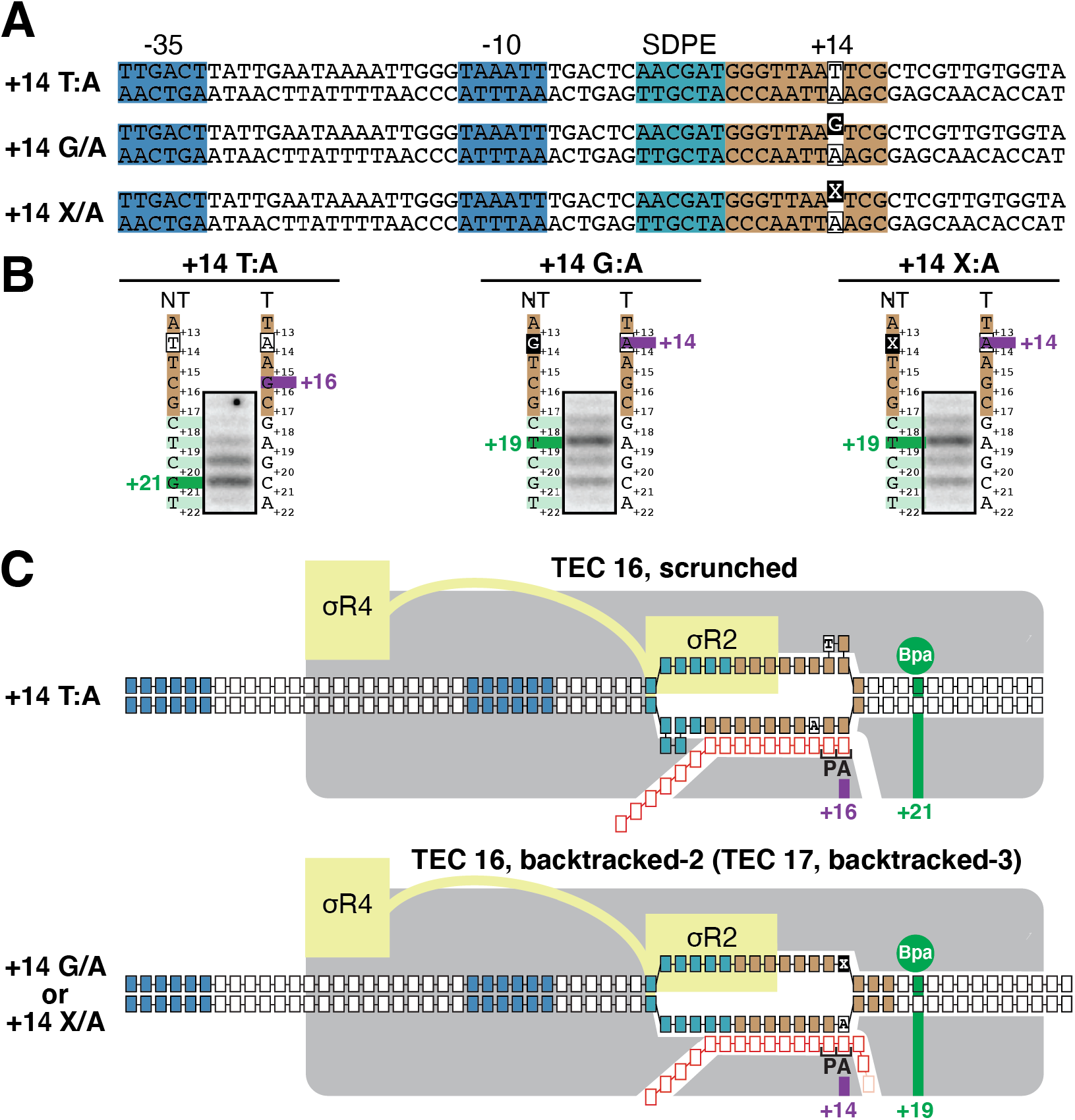
Strand-dependence of sequence determinants for scrunching in σ-dependent pausing. (A) *λ*PR’ promoter derivatives containing consensus nucleotide T at nontemplate strand position +14 (top row; +14 T:A), non-consensus nucleotide G at nontemplate strand position +14 (center row; +14 G/A), or abasic site at nontemplate strand position +14 (bottom row; +14 X/A). Raised black-filled box, non-consensus nucleotide or abasic site. Other colors as in Figure 1A. (B) Positions of RNAP leading edge on *λ*PR’ promoter derivatives of panel A. For each promoter derivative, figure shows gel image for primer-extension mapping of crosslinking sites, nontemplate- and template-strand sequences (to left and right of gel image; EPS colored as in Figure 1A), observed crosslinking sites (forest green), and inferred RNAP-active-center A-site position. (C) Mechanistic interpretation of data in panel B. Colors as in Figure 3B-C.

### Structural basis of scrunching in σ-dependent pausing

Structures of σ-containing TECs have been reported previously (18, 19), but a structure of a scrunched, paused, σ-containing transcription elongation complex, such as the species defined in results in Figures 3-5, has not been reported previously. To determine the structural basis of σ-dependent pausing, we performed cryo-EM structure determination, analyzing a σ-dependent pTEC prepared in solution. We incubated a synthetic nucleic-acid containing scaffold containing the *λ*PR’ promoter, a consensus SDPE positioned as in *λ*PR’, and a 15 bp non-complementary region, corresponding to the transcription bubble of the σ-dependent pTEC at *λ*PR’ (Figure 6A), with *E. coli* RNAP σ^70^ holoenzyme, and we applied samples to grids, flash-froze samples, and performed single-particle-reconstruction cryo-EM (Figures 6 and S6).

**Figure 6.**
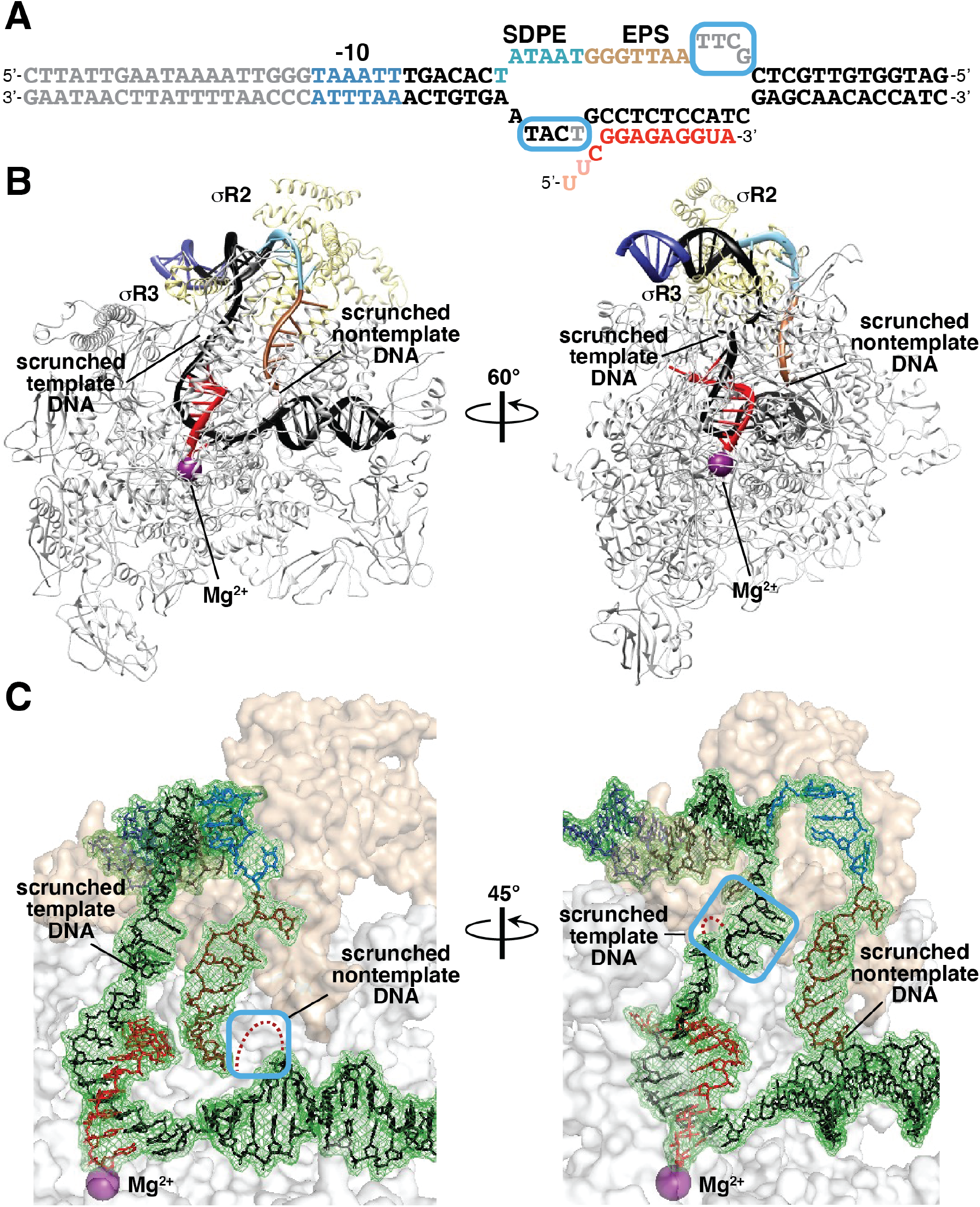
Structural basis for scrunching in σ-dependent pausing. (A) Nucleic-acid scaffold. DNA, black (−10 element, SDPE, EPS, and disordered nucleotides in blue, light blue, brown, and gray, respectively; non-complementary region corresponding to unwound transcription bubble indicated by raised and lowered letters); RNA, red (disordered nucleotides in pink); cyan boxes, nontemplate- and template-strand DNA nucleotides disordered or repositioned due to DNA scrunching. (B) Cryo-EM structure of scrunched σ-dependent paused TEC (pTEC; two orthogonal view orientations). Violet sphere, RNAP-active-center catalytic Mg^2+^. Other colors as in panel A. (C) Cryo-EM density and atomic model, showing interactions of RNAP and σ with DNA and RNA. Cyan boxes, nontemplate-strand (left) and template-strand (right) DNA nucleotides disordered or repositioned due to DNA scrunching; red dots, DNA nucleotides disordered due to DNA scrunching. Other colors as in panel A.

The cryo-EM structure of pTEC has an overall resolution of 3.8 Å (Figure S6). Map quality is high, with ordered, traceable, density for RNAP, σR2, σR3, 13 bp of upstream dsDNA, all except 4 nucleotides of the non-template strand of the transcription bubble, all except one nucleotide of the template strand of the transcription bubble, 13 bp of downstream dsDNA, and 12 nt of RNA, corresponding to the 12-nt segment closest to the RNA 3’ end (Figure 6B-C).

The cryo-EM structure of pTEC shows a σ-containing transcription elongation complex in an RNAP post-translocated state (Figure 6B-C). σR2 interacts with the SDPE in pTEC, making the same sequence-specific protein-DNA interactions as made by σR2 with the -10 element in RPo, and σR3 interacts with the DNA segment immediately upstream of the SDPE in pTEC, making the same protein-DNA interactions as made by σR3 in RPo (Figure 6B). The transcription-bubble length is 16 bp, corresponding to the 15 bp non-complementary region and one additional base pair immediately downstream of the non-complementary region (Figure 6B-C). The structure of pTEC shows 3 bp of DNA scrunching as compared to the structure of RPo (16 bp unwound vs. 13 bp unwound; (12, 73–75) and Figure S7A). Scrunching in pTEC results in disorder of 4 nontemplate-strand nucleotides at the downstream edge of the transcription bubble (*λ*PR’ positions +14 through +17; Figures 6C and S7A, left) and in disorder of one template-strand nucleotide and re-positioning of 3 template-strand nucleotides in the upstream part of the transcription bubble, between upstream double-stranded DNA and the RNA-DNA hybrid (*λ*PR’ positions +5 and +2 through +4; Figures 6C and S7A, right).

In RPo, σR2 having Bpa incorporated at position 448 crosslinks more strongly to the template-stand nucleotide at the third position of the -10 element than to the template-stand nucleotide at the fourth position of the -10 element (Figures 3A-B and S3), whereas, in pTEC, σR2 having Bpa incorporated at position 448 crosslinks less strongly to the template-stand nucleotide at the third position of the SDPE than to the template-stand nucleotide at the fourth position of the SDPE (Figure 3A,B). Comparison of the structures of RPo and pTEC explains this difference in σR2-DNA crosslinking in RPo and pTEC (Figure S7B). Namely, in RPo, residue 448 of σR2 is closer to the nucleotide at the third position of the -10 element (∼12 Å vs. ∼19 Å; Figure S7B, left), whereas, as a result of DNA scrunching and repositioning of template-strand nucleotides in pTEC, residue 448 of σR2 is closer to the fourth position of the SDPE (∼13 Å vs. ∼15 Å; Figure S7B, right).

The nontemplate strand of the nucleic-acid scaffold used to obtain the structure of pTEC contained a perfect match to the consensus sequence for formation of scrunched σ-dependent paused complexes and the consensus sequence for formation of scrunched initial-transcription paused complexes (positions +14 to +17; Figures 4C, top and middle, and 6A). The structure of pTEC shows that the nontemplate-strand nucleotide that is strongly specified in the consensus sequences for scrunched σ-dependent paused complexes and scrunched initial-transcription paused complexes (Figure 4C, top and middle)--but that is not strongly specified in the consensus sequence for elemental pausing in transcription elongation (Figure 4C, bottom)--is the first nucleotide of the 4-nucleotide nontemplate-strand segment that is disordered due to DNA scrunching (position +14; Figures 6B-C and S7A, left). The fact that this nucleotide is strongly specified in the consensus sequences for scrunched σ-dependent paused complexes and scrunched initial-transcription paused complexes (Figure 4C, top and middle)--but is not strongly specified in the consensus sequence for elemental pausing in transcription elongation, which does not involve DNA scrunching (Figure 4C, bottom; (76))--together with the observation that this nucleotide is the first nucleotide in the 4-nt nontemplate-strand segment re-positioned due to DNA scrunching, suggests that specificity at this position is associated with DNA scrunching and reflects sequence-dependent differences in the ability to accommodate DNA scrunching.

## Discussion

Our results: (1) establish, through mapping of positions of the RNAP leading and trailing edges relative to DNA, that σ-dependent pausing at *λ*PR’ involves a scrunched state with 2-3 bp of DNA scrunching and an unscrunched, backtracked state (Figure 3); (2) define a consensus sequence for formation and/or stability of a scrunched σ-dependent paused complex (T_P-1_N_P_Y_A_G_A+1_) that is identical to the consensus sequence for formation and/or stability of a scrunched initial-transcription paused complex (T_P-1_N_P_Y_A_G_A+1_) and similar to the consensus sequence for elemental pausing in transcription elongation (Y_A_G_A+1_) (Figure 4); (3) show the position that is strongly specified in the consensus sequences for the scrunched σ-dependent paused complex and the scrunched initial-transcription paused complex, but not in the consensus sequence for elemental pausing in transcription elongation (T_P-1_), is recognized through the nontemplate DNA strand (Figure 5); and (4) provide an atomic structure of a scrunched σ-dependent paused complex that suggests specificity at position T_P-1_ is associated with DNA scrunching and reflects sequence-dependent differences in the ability to accommodate DNA scrunching (Figure 6).

Our results provide dispositive evidence for the hypothesis that, in σ-dependent pausing at *λ*PR’, following pause capture, the pTEC extends RNA by 3-4 nt using a scrunching mechanism, and for the hypothesis that the resulting pTECs with extended RNA can collapse to yield a backtracked state (10, 41, 55–59).

DNA scrunching has been shown to mediate translocation of the RNAP active center relative to DNA in transcription-start-site selection, initial transcription, and promoter escape during transcription initiation (7-9, 63, 64, 68, 77). DNA stepping has been thought to mediate translocation of the RNAP active center relative to DNA during transcription elongation (29). Our results establish that this distinction between the mechanisms of RNAP-active-center translocation during transcription initiation and transcription elongation is not absolute, showing that DNA scrunching mediates translocation of the RNAP active center under certain circumstances in pausing and pause escape during transcription elongation. The shared feature of the transcription complexes that engage in DNA scrunching during transcription initiation and the transcription complexes shown here to engage in DNA scrunching in pausing and pause escape during transcription elongation is the presence of sequence-specific σ-DNA interactions that anchor the trailing edge of RNAP relative to DNA, preventing RNA extension through a DNA stepping mechanism, and thereby necessitating RNA extension through a DNA scrunching mechanism.

Generalizing from this observation, we propose that DNA scrunching occurs in pausing and pause escape during transcription elongation whenever sequence-specific σ-DNA interaction anchors the trailing edge of RNAP relative to DNA, including, for example, whenever a σ-containing TEC encounters an SDPE in a transcribed region (e.g., σ-dependent pausing at sequences other than *λ*PR’ SDPE; (41, 44–48, 54, 78, 79)), or whenever a sequence-specific DNA-binding protein that interacts with RNAP engages a σ-containing TEC in a transcribed region (e.g., transcription antitermination factor Q at a Q-binding element upstream of an SDPE (57), or a transcription activator protein, such as catabolite activator protein, CAP, able to interact with RNAP from an appropriately positioned DNA site upstream of an SDPE).

Generalizing further, we suggest that DNA scrunching occurs in pausing and pause escape during transcription elongation in *any* circumstance in which *any* sequence-specific protein-DNA interaction anchors the trailing edge of RNAP relative to DNA. Examples potentially include: (i) pausing induced by RfaH, which binds to a σ-free TEC and makes sequence-specific protein-DNA interactions with the transcription-bubble nontemplate-DNA strand similar to the sequence-specific interactions between σ and an SDPE (80, 81); (ii) pausing induced by other NusG/RfaH-family transcription factors (82), such as *Bacillus subtilis* NusG, that bind to a σ-free TEC and make sequence-specific protein-DNA interactions with the transcription-bubble nontemplate-DNA strand similar to the sequence-specific interactions between σ and an SDPE (83, 84); (iii) pausing induced by other factors that bind to a TEC and make sequence-specific protein-DNA interactions with an appropriately positioned upstream DNA site, and (iv) pausing induced by sequence-specific protein-DNA interaction between RNAP *α*subunit C-terminal domain (*α*-CTD) and an appropriately positioned upstream DNA site.

We suggest that the DNA scrunching that occurs in pausing and pause escape during transcription elongation--like the DNA scrunching that occurs in initial transcription and promoter escape during transcription initiation (1, 7–10)--serves as the mechanism to capture and store the free energy required to break the sequence-specific protein-DNA interactions that anchor RNAP on DNA. DNA scrunching during pausing enables capture of free energy from multiple nucleotide additions and stepwise storage of the captured free energy in the form of stepwise increases in the amount of DNA unwinding. Upon rewinding of the upstream part of the unwound DNA (e.g., the SDPE in σ-dependent pausing), the free energy captured and stored during scrunching is accessed to drive pause escape.

We note that the approaches of this work--specifically, mapping of positions of the RNAP leading and trailing edges relative to DNA using Bpa-modified RNAP--will enable direct determination whether, and if so how, DNA scrunching occurs in each of the above circumstances.

## Materials and Methods

Protein-DNA photocrosslinking was performed as in (63, 64). XACT-seq was performed as in (65), and the resulting data was analyzed using custom Python scripts. The cryo-EM structure was determined using single-particle reconstruction. Full details of methods are presented in SI Materials and Methods.

## Acknowledgments

Cryo-EM data were collected at the Rutgers University Cryo-EM and Nanoimaging Facility and the University of Michigan Life Sciences Institute Cryo-EM Facility. Cryo-EM data analysis were performed with assistance from Chengyuan Wang. We thank Jeremy Bird for providing GreB. Work was supported by National Institutes of Health grants ES030791 (MS), GM133777 (JBK), GM041376 (RHE), and GM118059 (BEN).

## SI Materials and Methods

### Strains

Plasmids were maintained in *E. coli* strain DH10B (Thermo Fisher Scientific) and NEB 5- alpha (New England Biolabs). Protein expression was performed using *E. coli* strain NiCo21(DE3) (New England Biolabs) or *E. coli* strain BL21 Star (DE3) (Invitrogen). *E. coli* strain ML149 (1) is a derivative of MG1655 that contains a kanamycin cassette downstream of *rpoD*. *E. coli* strain ML176 (1) is a *greA^-^* derivative of ML149. Strains ML149 and ML176 were used for analysis of σ-dependent pausing *in vivo*.

### Oligodeoxyribonucleotides

Oligonucleotides (Table S1) were dissolved in nuclease-free water to 1 mM and stored at -80°C.

### Plasmids

Plasmid pLHN12-His (2) contains the gene for *E. coli σ*^70^ with an N-terminal histidine coding sequence (6xHis) under the control of an isopropyl *β*-D-1-thiogalactopyranoside (IPTG)-inducible. Plasmid pLHN12-His-(R541C, L607P) is a derivative of pLHN12-His with substitutions that disrupt interaction between *σ*^70^ region 4 and the *β* flap (3).

Plasmid pIA900-*β′*^R1148Bpa^ and pIA900-*β′*^T48Bpa^ contains genes for the RNAP *β′* subunit with a nonsense codon (TAG) at position 1148 or 48, respectively, with an N-terminal decahistidine coding sequence (4).

Plasmid pEVOL-pBpF (Addgene; (5)) contains genes directing the synthesis of an engineered Bpa-specific UAG-suppressor tRNA and an engineered Bpa-specific aminoacyl-tRNA synthetase that charges the amber suppressor tRNA with Bpa.

Plasmid p*σ*^70^, is a derivative of pBAD24 ((6); American Type Culture Collection, ATCC), containing the gene for *E. coli σ*^70^ with an N-terminal decahistidine coding sequence under the control of an arabinose-inducible promoter (pBAD). To construct plasmid p*σ*^70^, we used PCR to generate a ∼4.6 kb DNA fragment containing a decahistidine coding sequence, pBAD promoter, *araC*, pBR322 *ori*, and ampicillin resistance gene and a ∼2 kb DNA fragment containing the gene for *E. coli σ*^70^. To generate the pBAD-containing fragment, reactions contained plasmid pBAD24 (0.04 μM), primer JW119 (0.5 μM), primer JW270 (0.5 μM), and 1X Phusion HF Master Mix (Thermo Fisher Scientific). To generate the *σ*^70^-containing fragment, reactions contained pET28-*rpoD* (0.04 μM; gift of J. Roberts), primer JW155 (0.5 μM), primer JW154 (0.5 μM), and 1X Phusion HF Master Mix. (30 cycles at 95°C, 10 sec; 55°C, 10 sec, 72°C, 4 min). After PCR, 20 U of DpnI (New England Biolabs) was added, reactions were incubated at 37°C for 16 hrs, and nondigested DNA fragments were isolated using a PCR purification kit (Qiagen). Recovered fragments (∼50 ng) were mixed with 1X Gibson assembly master mix (New England Biolabs) in a 20 μl reaction and incubated at 50°C for 20 min. Next, 1 μl of the Gibson assembly reaction was introduced by electroporation into DH10B cells (Invitrogen), cells were plated on LB agar containing 100 μg/ml carbenicillin, recombinant plasmid DNA was isolated from individual transformants, and plasmid sequences verified by Sanger sequencing (Macrogen/Psomagen).

Plasmid p*σ*^70^-R448Bpa-His^10^, is a derivative of plasmid p*σ*^70^ with a nonsense codon (TAG) at position 448 of the gene for *E. coli σ*^70^. Plasmid p*σ*^70^-R448Bpa-His^10^ was generated by site-directed mutagenesis. Reactions (12.5 μl volume) contained 0.04 μM of p*σ*^70^, 0.5 μM oligo HV75 and 1 X Phusion HF Master Mix (Thermo Fisher Scientific) (95°C for 2 min; 95°C for 15 sec, 55°C for 15 sec, 72°C for 2 min; 30 cycles). After PCR, 20 U of DpnI (New England Biolabs) was added, reactions were incubated at 37°C for 16 hrs. Next, 1 μl of reaction was introduced by electroporation into DH10B cells (Invitrogen), cells were plated on LB agar containing 100 μg/ml carbenicillin, recombinant plasmid DNA was isolated from individual transformants, and plasmid sequences verified by Sanger sequencing (Macrogen/Psomagen).

Plasmid pCDF-*λ*PR*′* is a derivative of pCDF-CP (7) that contains *λ*PR*′* sequences from positions -65 to +50 inserted into BglI-digested pCDF-CP. Plasmid pCDF-CP contains a CloDF13 replication origin, a selectable marker conferring resistance to spectinomycin and streptomycin, and two BglI recognition sites used to introduce DNA fragments upstream of transcription terminator tR2.

The +14-20 library was generated using procedures described in (8, 9) with JW615 as template and s1219 and s1220 as amplification primers. The amplification primers have 5*′* end sequences that introduce BglI recognition sequences and 3*′* end sequences complementary to the template oligo. PCR amplicons were treated with BglI and BglI-digested fragments were ligated into BglI-digested pCDF-CP. The ligation mixture was transformed into NEB 5-alpha cells (New England Biolabs), cells plated on LB agar plates containing 50 μg/ml spectinomycin and 50 μg/ml streptomycin, and recombinant plasmid DNA (pCDF-*λ*PR*′*-N7) was isolated from *∼*0.5 x 10^6^ transformants.

### Proteins

RNAP core enzyme was prepared from *E. coli* strain NiCo21(DE3) (New England Biolabs) containing plasmid pIA900 (10) using procedures described in (4).

Bpa-containing RNAP core enzyme derivatives, RNAP-*β′*^R1148Bpa^ and RNAP-*β′*^T48Bpa^, were prepared from *E. coli* strain NiCo21(DE3) (New England Biolabs) containing plasmid pIA900-RNAP-*β′*^R1148Bpa^ or plasmid pIA900-RNAP-*β′*^T48Bpa^ (4) and plasmid pEVOL-pBpF (5), using procedures described in (4).

Wild-type *σ*^70^ was purified using procedures described in (11). Bpa-containing *σ*^70^ derivative, *σ*^70 R448Bpa^, was prepared from *E. coli* strain DH10B (Invitrogen) containing plasmid p*σ*^70^-R448Bpa-His^10^ using procedures described in (4). *σ*^70 C541, P607^ was prepared from *E. coli* strain BL21 Star (DE3) (Invitrogen) containing pLHN12-His-(R541C, L607P) using procedures described in (2).

RNAP *σ*^70^ holoenzyme was prepared by incubating 1 μM *E. coli* RNAP core enzyme and 5 μM *E. coli σ*^70^ in 10 mM Tris-Cl (pH 8.0), 100 mM KCl, 10 mM MgCl_2_, 0.1 mM EDTA, 1 mM DTT, and 50% glycerol for 30 min at 25°C.

GreB was purified using procedures described in (12).

### DNA templates for *in vitro* assays

DNA templates used for *in vitro* assays in Figures 1C, 3A, and S1-S4, which contain sequences from positions -86 to +96 of *λ*PR*′* promoter, were generated by PCR amplification of *∼*1 pg pCDF-*λ*PR*′*with 0.4 μM primer JW521 and 0.4 μM primer JW544 in reactions containing 1 X Phusion HF Master Mix (Thermo Fisher Scientific). Amplicons were purified using a PCR purification kit (Qiagen).

DNA templates used for *in vitro* assays in Figure 5, which contain sequences from positions -65 to +79 of *λ*PR*′* promoter with a T, G or X (where X is an abasic site) at nontemplate strand position +14, were prepared by mixing 0.4 μM of template-strand oligo (JW680) with 0.4 μM nontemplate strand oligo (JW679, JW683 or JW687) in 1 X Phusion HF Master Mix (Thermo Fisher Scientific) and performing 30 cycles of 95°C for 2 min; 95°C for 15 sec, 55°C for 15 sec, and 72°C for 2 min. Double stranded products were gel purified to remove non-annealed oligos.

### σ-dependent pausing *in vitro*

*In vitro* transcription assays in Figure 1C and S1 were performed essentially as described in (3, 13). Reactions in Figure 1C (120 μL total volume) contained 10 nM of *λ*PR*′* template, 40 nM of RNAP holoenzyme (wild-type), 200 μM each of ATP, GTP, CTP and UTP supplemented with 0.03 mCi *α*-^32^P-UTP [Perkin Elmer; 3000 Ci/mmol]) in 1X reaction buffer (10 mM Tris-HCl, pH 8.0; 70 mM NaCl, 0.1 mg/ml bovine serum albumin, BSA; and 5% glycerol) and 100 nM GreB (where indicated) were incubated for 10 min at 37°C. Next, a mixture of 10 mM MgCl_2_ and 10 μg/ml rifampin (Gold Biotech) was added, 20 μL aliquots were removed at the indicated times (20s, 40s, 60s, 80s), and mixed with five volumes of stop solution (500 mM Tris-HCl, pH 8.0; 10 mM EDTA; and 0.1 mg/ml glycogen). 120 μL of phenol:chloroform (pH 4.5; Ambion) was added, samples were mixed, the aqueous layer (∼100 μL) was removed, 100% ethanol was added (∼300 μL), samples were placed at −80°C for 12 h, precipitated nucleic acids were recovered by centrifugation, re-suspended in 5 μl nuclease free water, and mixed with 5 μL loading dye (1 X TBE, 8 M urea, 0.025% xylene cyanol, and 0.025% bromophenol blue). Samples were heated at 95°C for 2 min, cooled to 95°C, and analyzed by electrophoresis on 20%, 8 M urea, 1 X TBE polyacrylamide gels (Urea Gel System; National Diagnostics). Radiolabeled bands were visualized by storage phosphor screen (GE Healthcare) and phosphorimagery (Typhoon 9400 variable mode imager, GE Healthcare). Sizes of RNA products were estimated by comparison to radiolabeled Decade Marker (Thermo Fisher Scientific).

Reactions in Figure S1 (20 μL total volume) contained 4 nM of the *λ*PR*′*linear DNA template, 40 nM of the indicated RNAP holoenzyme, 1X RB (10 mM Tris–HCl, pH 8.0; 70 mM NaCl; 10 mM MgCl_2_; 0.1 mg/ml BSA; and 5% glycerol) and 100 nM GreB (where indicated) were incubated for 10 min at 37°C, 200 μM ATP, 200 μM GTP, 200 μM CTP and 200 μM UTP supplemented with 0.02 mCi *α*-^32^P-UTP (Perkin Elmer; 3000 Ci/mmol) were added, reactions were incubated at 37°C for 10 min, 100 μL of stop solution was added, and samples further processed as described in the preceding paragraph.

### σ-dependent pausing *in vivo*

ML149 (*greA*^+^) and ML176 (*greA^-^*) cells containing plasmid pCDF-*λ*PR*′* were grown in 25 ml LB containing 100 μg/ml carbenicillin and 1 mM IPTG in 125 ml DeLong flasks (Bellco Glass), shaken at 37°C on an orbital platform shaker at 220 rpm. When cultures reached an OD_600_ of ∼0.5, 1 ml aliquots of cell suspensions were removed before or at the indicated time points after addition of 500 μg/ml rifampin (Gold Biotech). 1 ml aliquots of cell suspensions were mixed with 3 ml of RNAlater solution (Thermo Fisher Scientific) in 50 ml Oakridge tubes by inversion several times and incubated overnight at 4°C. The mixture was centrifuged at 17,000 × g for 20 min at 4°C, supernatant was removed, 1 ml of Tri-reagent (Molecular Research Center) was added and pellets were dispersed by vortexing. Cell suspensions in Tri-reagent were transferred to 1.7 ml low binding tubes (Axygen), incubated at 70°C for 10 min, centrifuged at 21,000 × g at 4°C for 10 min, and the supernatants were recovered into fresh tubes. 200 μl of chloroform was added to each tube and mixed by vigorous shaking for 15 s. Phases were separated by centrifugation at 21,000 × g at 4°C for 15 min. 500 μl of the upper, aqueous phase was recovered and transferred to a fresh tube to which 167 μl of 100% ethanol was added. Subsequent removal of RNA >200 nt and recovery of RNA <200 nt was performed using the mirVana microRNA Isolation kit (Thermo Fisher Scientific) according to the manufacturer’s protocol. After elution from mirVana columns, eluates were concentrated by ethanol precipitation and resuspended directly into formamide loading dye (95% deionized formamide, 18 mM EDTA, and 0.025% SDS, xylene cyanol, bromophenol blue, amaranth).

50 pmol of a locked nucleic acid (LNA) probe complementary to PR’ sequences +1 to +17 (LNA8) was incubated in a 20 μl volume with 5 μl *γ*-^32^P-ATP (EasyTide; Perkin Elmer), 2 μl 10X T4 PNK buffer, 9 μl nuclease free water (Thermo Fisher Scientific), and 2 μl T4 PNK (New England Biolabs) at 37°C for 1 hr followed by 95°C for 10 min. Labeled probe was separated from unincorporated radiolabeled nucleotide using a size-exclusion spin column (Cytiva Illustra Microspin G-25; Thermo Fisher Scientific).

Pause RNAs and full-length RNAs generated from *λ*PR*′ in vivo* were detected by hybridization as described in (14, 15). RNA isolated from cells was subjected to electrophoresis on 20% 8M urea slab gels (equilibrated and run in 50 mM MOPS, pH 7), transferred to a neutral nylon membrane (Whatman Nytran N; GE Healthcare Life Sciences) using a semi-dry electroblotting apparatus (Biorad) operating at 20V for 25 min using chilled 20 mM MOPS (pH 7) as conductive medium. RNA was crosslinked to the membrane using 157 mM

N-(3-dimethylaminopropyl)-N′-ethylcarbodiimide hydrochloride (EDC) (Sigma–Aldrich) in 0.97% 1-methylimidazole (pH 8) (Alfa Aesar) for 60 min at 55°C. Membrane was washed with 20 mM MOPS (pH 7) at 25°C, placed on nylon hybridization mesh, the membrane mesh stack was placed into a hybridization bottle (70 × 150 mm) at 50°C and 50 ml of pre-hybridization solution (5X SSC, 5% SDS, 2X Denhardt’s solution, 40 μg/ml sheared salmon sperm DNA solution [Thermo Fisher Scientific], 20 mM Na_2_HPO_4_ [pH 7.2] in diethylpyrocarbonate (DEPC) treated water) at 50°C was added. The hybridization bottle was incubated at 50°C for 30 min with constant rotation, the solution was decanted and replaced by a 50 ml of pre-warmed hybridization solution with radiolabeled LNA probe prepared above and continue incubation at 50°C for 16 hr. The membrane was washed in non-stringent wash buffer (3X SSC, 5% SDS, 10X Denhardt’s solution, 20 mM Na_2_HPO_4_ [pH 7.2] in DEPC treated water) two times for 10 min, two times for 30 min and in stringent wash buffer (1X SSC, 1% SDS, in DEPC treated water) one time for 5 min before it was blotted dry, wrapped in plastic film, and radiolabeled bands were visualized by storage phosphor screen (GE Healthcare) and phosphorimagery (Typhoon 9400 variable mode imager, GE Healthcare).

### *In vitro* protein-DNA photocrosslinking

*In vitro* protein-DNA photocrosslinking and crosslink mapping experiments were performed as described in (7, 16). Reactions contained 40 nM RNAP holoenzyme (*β′*T48Bpa, *σ*^70^R448Bpa or *β′*R1148Bpa), 4 nM template, 1 X RB (10 mM Tris-Cl, pH 8.0; 70 mM NaCl; 10 mM MgCl_2_; 0.1 mg/ml BSA; and 5% glycerol) and 100 nM GreB (where indicated) were incubated for 2 min at 37°C, 200 μM ATP, 200 μM CTP, 200 μM GTP, and 200 μM UTP were added, reactions were further incubated for 10 min, and subjected to UV irradiation for 10 min at 25°C in a Rayonet RPR-100 photochemical reactor equipped with 16 x 350 nm tubes (Southern New England Ultraviolet).

Reactions were mixed with 15 μL 5 M NaCl and 6 μL 100 μg/μl heparin, incubated for 5 min at 95°C and then cooled to 4°C. RNAP-DNA crosslinked complexes were isolated by adding 20 μl MagneHis Ni-particles (Promega) equilibrated and suspended in 1 X Taq DNA polymerase buffer, 10 mg/ml heparin, and 0.1 mg/ml BSA; MagneHis Ni-particles were collected using a magnetic microfuge tube rack; particles were washed with 1 X Taq DNA polymerase buffer, 10 mg/ml heparin, and 0.1 mg/ml BSA, washed twice with 50 μl 1 X Taq DNA polymerase buffer (New England Biolabs), and particles (which contained bound RNAP-DNA complexes) were resuspended in 10 μl 1 X Taq DNA polymerase buffer.

Primer extension reactions (12.5 μl) were performed by combining 2 μl of the recovered RNAP-DNA complexes, 1 μl of 1 μM ^32^P-5*′* end-labeled primer JW77 (for leading edge position) or JW544 (for trailing edge and *σ*R2), 1 μL 10 X dNTPs (2.5 mM dATP, 2.5 mM dCTP, 2.5 mM dGTP, 2.5 mM dTTP), 0.25 μL 5 U/ml Taq DNA polymerase (New England Biolabs), 5 μL 5 M betaine, 0.625 μL 100% dimethyl sulfoxide, and 1.25 μl 10 X Taq DNA polymerase buffer; 40 cycles of 30 sec at 95°C, 30 sec at 55°C, and 30 sec at 72°C. Reactions were stopped by addition of 12.5 μL 1 X TBE, 8 M urea, 0.025% xylene cyanol, and 0.025% bromophenol blue. Radiolabeled products were separated by electrophoresis on 8% 8M urea slab gels (equilibrated and run in 1 X TBE) and visualized by storage-phosphor imaging (Typhoon 9400 variable-mode imager; GE Life Science). Positions of RNAP-DNA crosslinks were determined by comparison to products of a DNA-nucleotide sequencing reaction generated using oligo JW77 or JW544 and a DNA template containing sequences from positions -86 to +96 of pCDF-*λ*PR*′* (Thermo Sequenase Cycle Sequencing Kit; Affymetrix).

### *In vivo* protein-DNA photocrosslinking

*In vivo* protein-DNA photocrosslinking and crosslink mapping experiments were done essentially as in (16, 17). Experiments in Figure 3A and S2-S4 were performed by sequential introduction of plasmid pCDF-*λ*PR*′*, plasmid pIA900-RNAP-*β′*^T48Bpa^ or pIA900-RNAP-*β′*^R1148Bpa^ or p*σ*^70 R448Bpa^ and plasmid pEVOL-pBpF into electrocompetent *E. coli* strain NiCo21(DE3) by transformation. After the final transformation step, cells were plated on LB agar containing 100 μg/ml carbenicillin, 50 μg/ml spectinomycin, 50 μg/ml streptomycin, and 25 μg/ml chloramphenicol; at least 1,000 individual colonies were scraped from the plate, combined, and used to inoculate 250 ml LB broth containing 1 mM Bpa (Bachem), 100 μg/ml carbenicillin, 50 μg/ml spectinomycin, 50 μg/ml streptomycin, and 25 μg/ml chloramphenicol in a 1000 mL flask (Bellco Glass) to yield OD_600_ = 0.3; the culture was placed in the dark and shaken (220 rpm) for 1 h at 37°C; 1 mM IPTG (for pIA900-RNAP-*β′*^R1148Bpa^ or pIA900-RNAP-*β′*^T48Bpa^) or 0.2% L-arabinose (for *σ*^70 R448Bpa^) was added to induce expression; and the culture was placed in the dark and shaken (220 rpm) for 3 hrs at 37°C (for pIA900-RNAP-*β′*^R1148Bpa^ or pIA900-RNAP-*β′*^T48Bpa^ expression) or 3 hrs at 30°C (for *σ*^70 R448Bpa^ expression).

UV irradiation of cell suspension, purification of RNAP-DNA photocrosslinked complexes from cell cultures, denaturation, isolation, primer extension and electrophoresis for mapping RNAP-DNA crosslinked complexes were done following the procedures as described in (16, 17).

Analysis of *σ*-dependent pausing *in vivo* for +14-20 library transcription complexes (Figure 4) was performed by sequential introduction of plasmid pCDF-*λ*PR*′*-N7 (+14-20) library (yielding *∼*28 million transformants), plasmid pIA900-RNAP-*β*’^R1148Bpa^ (yielding *∼*8 million transformants), and plasmid pEVOL-pBpF (yielding *∼*5 million transformants) into electrocompetent *E. coli* strain NiCo21(DE3). After the final transformation step, cells were plated on *∼*4-6 LB agar plates containing 100 μg/ml carbenicillin, 50 μg/ml spectinomycin, 50 μg/ml streptomycin, and 25 μg/ml chloramphenicol to yield a lawn. Colonies were scraped from the surface of the plates, combined, and used to inoculate 150 ml LB broth containing 1mM Bpa, 100 μg/ml carbenicillin, 50 μg/ml spectinomycin, 50 μg/ml streptomycin, and 25 μg/ml chloramphenicol in a 1000 mL flask to yield OD_600_ = 0.3; the culture was placed in the dark and shaken (220 rpm) for 1 h at 37°C; IPTG was added to 1 mM and the culture was placed in the dark and shaken (220 rpm) for 3 h at 37°C.

To measure background signal, a portion of the cell cultures containing pCDF-*λ*PR*′* or pCDF-*λ*PR*′*-N7 (+14-20 library) were removed, rifampin (Gold Biotech) was added to a final concentration of 200 μg/ml, and the culture was shaken at 37°C for 10 min prior to UV irradiation.

UV irradiation of cell suspension, purification of RNAP-DNA photocrosslinked complexes from cell cultures, denaturation, isolation, primer extension and electrophoresis for mapping RNAP-DNA crosslinked complexes were done following the procedures as described in (16, 17).

XACT-seq experiments (see below) were performed using denatured RNAP-DNA complexes isolated from cells containing the +14-20 library.

### XACT-seq: primer extension

Primer extension was performed in 50 µl reactions containing 8 µl of recovered RNAP-DNA complexes, 1 µl of 10 µM primer s128a, 5 μl 10 X dNTPs (2.5 mM dATP, 2.5 mM dCTP, 2.5 mM dGTP, 2.5 mM dTTP), 1 μl 5 U/μl *Taq* DNA polymerase, 20 μl 5 M betaine, 2.5 μl 100% dimethyl sulfoxide, and 5 µl 10 X *Taq* DNA polymerase buffer, and cycling 40 times through 30 s at 95°C, 30 s at 55°C, and 30 s at 72°C. Primer extension products were isolated by phenol:chloroform:IAA pH 8.0 extraction followed by ethanol precipitation, washed twice with 80% cold ethanol, resuspended in 20 µl water, and mixed with 20 µl of 2 X RNA loading dye (95% deionized formamide, 18 mM EDTA, 0.25% SDS, xylene cyanol, bromophenol blue, amaranth).

Primer extension products were separated by electrophoresis on 10% 7M urea slab gels (equilibrated and run in 1 X TBE), stained with SYBR Gold nucleic acid gel stain (Life Technologies) and ssDNA products ∼50- to ∼100-nt in size were excised from the gel. The gel fragment was crushed to elute nucleic acid from gel as described in (18), 350 μl of 0.3 M NaCl in 1 X TE buffer was added, the mixture was incubated for 10 min at 70°C, and the supernatant was collected using a Spin-X column (Corning). The elution procedure was repeated, supernatants were combined, and nucleic acids were recovered by ethanol precipitation, washed twice with 80% cold ethanol, and resuspended in 5 μl of nuclease-free water.

### XACT-seq: 3′-adapter ligation and library amplification

The recovered primer extension products (5 μl) were combined with 1 μl 10 X NEBuffer 1, ∼0.8 μM 3′-adapter oligo s1248 [5′ adenylated and 3′-end blocked oligo containing ten randomized nucleotides (10N) at the 5′ end], 5 mM MnCl_2_ and 1 μM of 5′-AppDNA/RNA ligase (New England Biolabs) in a final volume of 10 μl. The mixture was incubated for 1 h at 65°C followed by 3 min at 90°C, and cooled to 4°C for 5 min. The reaction was combined with 15 μl of mixture containing 10 U of T4 RNA ligase 1 (New England Biolabs), 1 X T4 RNA ligase 1 reaction buffer, 12% PEG 8000, 10 mM DTT, 60 μg/mL BSA. Reactions were incubated at 16°C for 16 h.

Adapter-ligated products were separated by electrophoresis on 10% 7M urea slab gels (equilibrated and run in 1 X TBE), stained with SYBR Gold nucleic acid gel stain and species ranging from ∼80 to ∼150-nt (for reactions containing oligo s1248 and primer extension products) or ∼50 and 90 nt (for reactions containing oligo s1248 and oligo JW402) were isolated by gel excision. The gel fragment was crushed, 400 μl of 0.3M NaCl in 1 X TE buffer was added, the mixture was incubated for 2 h at 37°C, the supernatant was collected using a Spin-X column (Corning). The elution procedure was repeated, supernatants were combined, and nucleic acids were recovered by ethanol precipitation, washed twice with 80% cold ethanol, and resuspended in 13 μl of nuclease-free water.

Adapter-ligated DNA (1 μl) were used as template in emulsion PCR (ePCR). Reactions contained 1 X Detergent-free Phusion HF reaction buffer containing 5 μg/ml BSA, 0.4 mM dNTPs, 0.5 μM Illumina RP1 primer, 0.5 μM Illumina index primer and 0.04 U/μl Phusion HF polymerase [95°C for 10 s, 95°C for 5 s, 60°C for 5 s, 72°C for 15 s (20 cycles), 72°C for 5 min]. Amplicons were recovered using a Micellula DNA Emulsion and Purification Kit. The emulsion was broken, DNA was purified according to the manufacturer’s recommendations, recovered by ethanol precipitation, and resuspended in 15 μl of nuclease-free water. Reaction products were separated by electrophoresis on a non-denaturing 10% slab gel (equilibrated and run in 1 X TBE), and amplicons between ∼150 bp and ∼220 bp were isolated by gel excision. The gel fragment was crushed, 400 μl of 0.3M NaCl in 1 X TE buffer was added, the mixture was incubated for 2 h at 37°C, the supernatant was collected using a Spin-X column. The elution procedure was repeated, supernatants were combined, and nucleic acids were recovered by isopropanol precipitation, washed twice with 80% cold ethanol, and resuspended in 15 μl of nuclease-free water.

Libraries generated by this procedure are: CP49, CP50, CP51, CP52, CP53 and CP54.

### XACT-seq: analysis of template sequences in the +14-20 library

To identify template sequences present in the +14-20 library, we performed ePCR in reactions containing ∼10^9^ molecules of the pCDF-*λ*PR*′*-N7 (+14-20) plasmid library, 1 X Detergent-free Phusion HF reaction buffer with 5 μg/ml BSA, 0.4 mM dNTPs, 0.5 μM Illumina RP1 primer, 0.5 μM Illumina index primer and 0.04 U/μl Phusion HF polymerase [95°C for 10 s, 95°C for 5 s, 60°C for 5 s, 72°C for 15 s (30 cycles), 72°C for 5 min]. Amplicons were recovered using a Micellula DNA Emulsion and Purification Kit. The emulsion was broken, DNA was purified according to the manufacturer’s recommendations, recovered by ethanol precipitation, and resuspended in 15 μl of nuclease-free water. Products were subjected to electrophoresis on a non-denaturing 10% slab gel (equilibrated and run in 1 X TBE), the 251 bp fragment was excised from the gel. The gel fragment was crushed, 400 μl of 0.3M NaCl in 1 X TE buffer was added, the mixture was incubated for 2 h at 37°C, the supernatant was collected using a Spin-X column. The elution procedure was repeated, supernatants were combined, and nucleic acids were recovered by isopropanol precipitation, washed twice with 80% cold ethanol, and resuspended in 15 μl of nuclease-free water. The library generated by this procedure is CP61T.

### XACT-seq: high-throughput sequencing

Barcoded libraries were pooled and sequenced on an Illumina NextSeq500 platform in high-output mode using custom sequencing primer s1115.

### XACT-seq: sample serial numbers

CP52, CP53 and CP54 are samples used for the identification of RNAP-active-center A-site positions in active transcription complexes in vivo (- Rifampin). CP49, CP50 and CP51 are samples used for the identification of RNAP-active-center A-site positions in static transcription complexes in vivo (+ Rifampin).

Sample CP61T was used to identify template sequences present in the +14-20 library.

### XACT-seq data analysis: ligation and crosslinking bias correction

To quantify crosslinking and ligation bias in the XACT-seq protocol, we reanalyzed results of (16). In that study, XACT-seq was performed in the presence and absence of rifampin (Rif+ and Rif-, respectively) on a p*lac*CONS promoter library that contained an 11 bp variable region (*lac*VR) spanning positions +3 to +13 relative to the transcription start site (TSS).

We obtained Illumina read counts from three Rif+ samples (CP22, CP24, and CP28). For each 11-nt *lac*VR sequence 𝑠, each crosslinked position 𝑝 within the larger p*lac*CONS promoter (defined as the 3’-most nucleotide of the primer extension product, with 𝑝 = 1 denoting the first position of the *lac*VR), and each sample X, we denote this quantity by 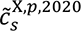. We also obtained corresponding Illumina read counts, 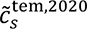, representing template abundance, i.e., the relative abundance of each DNA sequence 𝑠 in the plasmid library (sample CP26T). From these counts we computed the marginal nucleotide counts of each base 𝑏 = A, C, G, T at each position 𝑞 = 1, … ,11 within the *lac*VR:

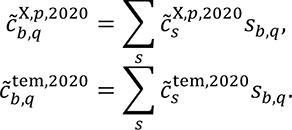

Here, 𝑠_b,q_ is a one-hot encoding of sequence 𝑠, i.e., 𝑠_b,q_ = 1 if base 𝑏 occurs at position 𝑞 in sequence 𝑠, and 𝑠_*,+_ = 0 otherwise.

From these counts we computed a weight matrix that quantifies the relative log_2_ probability of crosslinking and ligation based on the DNA sequence in the vicinity of the crosslink position. Specifically, let 𝑖 = 𝑞 − 𝑝 denote the relative coordinate of position 𝑞 in the *lac*VR with respect to the crosslink position 𝑝. The elements 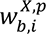 of this weight matrix were computed as

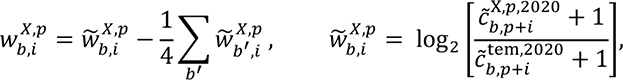

We restricted our attention to crosslink positions associated with the rifampin-inhibited initiation complex, namely, positions 𝑝 = +7 and +8 (which correspond to RNAP-active-center A- site positions +2 and +3, respectively). We reasoned that any sequence-dependence observed in our data at 𝑝 = +7 and +8 in the rifampin-inhibited complex would primarily reflect crosslinking and ligation bias.

At 𝑝 = +7 and +8, we found that the relative nucleotide positions 𝑖 = −2, −1,0, +1, +2 contributed substantially and reproducibly to the weight matrix elements 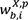, whereas other relative nucleotide positions 𝑖 did not. We therefore restricted the bias weight matrix to this range of values for 𝑖. We then averaged together the six matrices (corresponding to 𝑝 = +7, +8, and X = CP22, CP24, CP28) to generate a weight matrix quantifying crosslink and ligation bias:

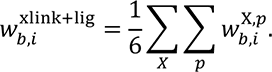

### XACT-seq data analysis: template bias correction

To correct for template bias, we obtained Illumina read counts, 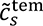, for the underlying plasmid library (sample CP61T). We then computed the marginal base counts at each nucleotide position 𝑞 = 1, … ,7 of the λPR’ variable region (λVR; promoter positions +14 to +20):

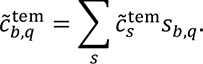

These counts were converted to a weight matrix that quantifies the relative log_2_ probability of each base at each position within the λVR:

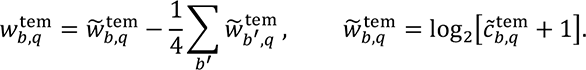

### XACT-seq data analysis: computation of reweighted sequence counts

To correct for bias in our XACT-seq data (both “xlink+lig” and “tem”), we reweighted the counts 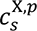 for each sequence 𝑠, sample 𝑋, and crosslink position 𝑝 (with position 𝑝 = 1 corresponding to the first position of λVR) using,

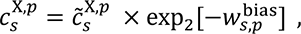

where

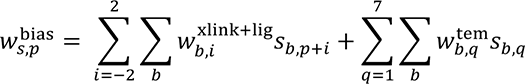

is the relative log_2_ probability of a sequence 𝑠 being both present in the variant λPR’ plasmid library and successfully crosslinked and ligated at position 𝑝 in the corresponding rifampin- inhibited initiation complex.

A similar correction was also performed for XACT-seq data reported in (16) that was obtained in the absence of rifampin.

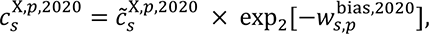

where

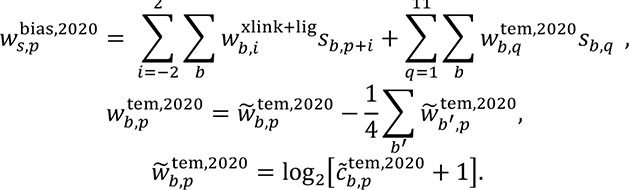

### XACT-seq data analysis: sequence logo generation

All sequence logos in this manuscript were created using Logomaker (19) and quantify the relative log_2_ enrichment of each possible base at each position within a sequence. Specifically, to compute a sequence logo from the (potentially reweighted) counts 𝑐_S_ of a set of sequence 𝑠, we first computed the marginal base counts at each position 𝑞 within the variable region,

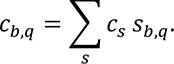

A logo was then generated in which the height ℎ_b,q_ of each base 𝑏 at each position 𝑞 within the logo was given by the centered log_2_ ratio

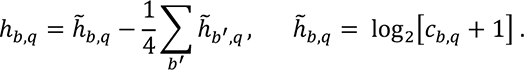

The logo for the σ-dependent paused complex (Figure 4C, top) was created by first generating three different logos, corresponding to position 𝑝 = +3 (A site position +16 relative to the TSS) and Rif- samples X = CP52, CP53, CP54. The logo shown was created by averaging these heights ℎ_b,q_ (for 𝑞 = 1,2,3,4) across the three replicates. Fig. S5B displays the same information but for all seven randomized positions (𝑞 = 1, … ,7). The logo for the initial-transcription paused complex (Figure 4C, center) was created by first generating three different logos, corresponding to position 𝑝 = +4 (A site position +6 relative to the TSS) and Rif- samples X = CP21, CP23, CP27. The logo shown was created by averaging these heights ℎ_b,q_ (for 𝑞 = 2,3,4,5) across the three replicates. The logo for the elemental paused complex (Figure 4C, bottom), was created from pause sites identified by NET-seq and reported in (20), with each sequence 𝑠 assigned a count 𝑐_s_= 1 and the position of the A site corresponding to the RNA 3’ end.

### Cryo-EM structure determination: sample preparation

The *σ*-dependent paused transcription elongation complex was prepared by incubating 12 μM *E. coli* RNAP *σ*^70^ core enzyme with 22 μM nucleic-acid scaffold (Figure 6) in 60 μl transcription buffer (20 mM Tris-HCl, pH 7.9, 30 mM KCl, 10 mM MgCl_2_, and 1 mM dithiothreitol) for 15 min at 25°C, followed by adding 20 μl 120 μM *E. coli* [C541; P607]-*σ*^70^ and incubating 10 min at 25°C. The sample was concentrated to 50 μl using 0.5 ml Amicon Ultra 30 kDa MWCO centrifugal concentrator (Millipore), mixed with 6 μl 80 mM CHAPSO, and stored on ice prior to applying on grids. 3 μl samples were applied to QuantiFoil 1.2/1.3 Cu 300-mesh grids (glow-discharged for 50 s) using a Vitrobot Mark IV autoplunger (FEI), with the environmental chamber set at 22°C and 100% relative humidity. Grids were blotted with filter discs #595 (Ted Pella) for 8 sec at blotting force 4, and flash-frozen in liquid ethane cooled with liquid nitrogen; grids were stored in liquid nitrogen.

### Cryo-EM structure determination: data collection and data processing

Cryo-EM data were collected at the University of Michigan Cryo-Electron Microscopy Facility, using a 300 kV Titan Krios G4i (Thermo Fisher Scientific) electron microscope equipped with a K3 direct electron detector and BioQuantum energy filter (Gatan). Data were collected automatically in counting mode using Leginon (21), a nominal magnification of 105,000x in EFTEM mode (actual magnification 595,238x), a calibrated pixel size of 0.84 Å per pixel, and a dose rate of 15 electrons/pixel/s. Movies were recorded at 100 ms/frame for 3 s (30 frames), resulting in a total radiation dose of 45 electrons/Å^2^. Defocus range varied between -1.5 μm and -3.5 μm. A total of 11,611 micrographs were recorded from one grid over five days. Micrographs were saved in Tiff format upon pre-processing for gain normalization and defect correction.

Data were processed as in Figure S6. Dose weighting and motion correction (5×5 tiles; b-factor = 150) were performed using Motioncor2 (22). CTF estimation was performed using CTFFIND4 (23). Subsequent image processing was performed using cryoSPARC (24). Automatic particle picking with blob picker yielded an initial set of 2,156,960 particles. Particles were binned 2x, extracted into 192 x 192-pixel boxes, and subjected to four rounds of reference-free 2D classification and removal of poorly populated classes, yielding a selected set of 921,867 particles. *Ab initio* models were generated, and heterogeneous refinement was performed for further classification of particles. One class, comprising particles with intact transcription complexes, was selected, and was subjected to homogeneous refinement, yielding a reconstruction with a global resolution of 3.8 Å as determined from gold-standard Fourier shell correlation.

The initial atomic model for protein components of the *σ*-dependent paused transcription elongation complex was built by manual docking of a cryo-EM structure of the *E. coli* Q21 transcription anti-termination loading complex Q21-QBE (PDB 6P18; (25)), with Q21 omitted, to the map in Chimera (26). Initial atomic models for DNA and RNA of the *σ*-dependent paused transcription elongation complex were built manually using Coot (27). The initial model of the of the *σ*-dependent paused transcription elongation complex was real-space rigid-body refined in Phenix (28) and subsequently refined with secondary-structure, geometry, Ramachandran, rotamer, Cβ, non-crystallographic-symmetry, and reference-model restraints. Molecular graphics representations were created using PyMOL and Chimera. EM density maps were visualized using PyMOL and Chimera.

### Data and software availability

Sequencing reads have been deposited in the NIH/NCBI Sequence Read Archive under the study accession number PRJNA797396. Source code and documentation are provided at http://www.github.com/jbkinney/21_nickels. A snapshot of this repository will be deposited on Zenodo prior to publication.

The final atomic model and map of the *σ*-dependent paused transcription elongation complex have been deposited in the Protein Data Bank (PDB) and Electron Microscopy Data Bank (EMD) with accession codes 7N4E and EMD-24148, respectively.

## SI Figures

**Figure S1.**
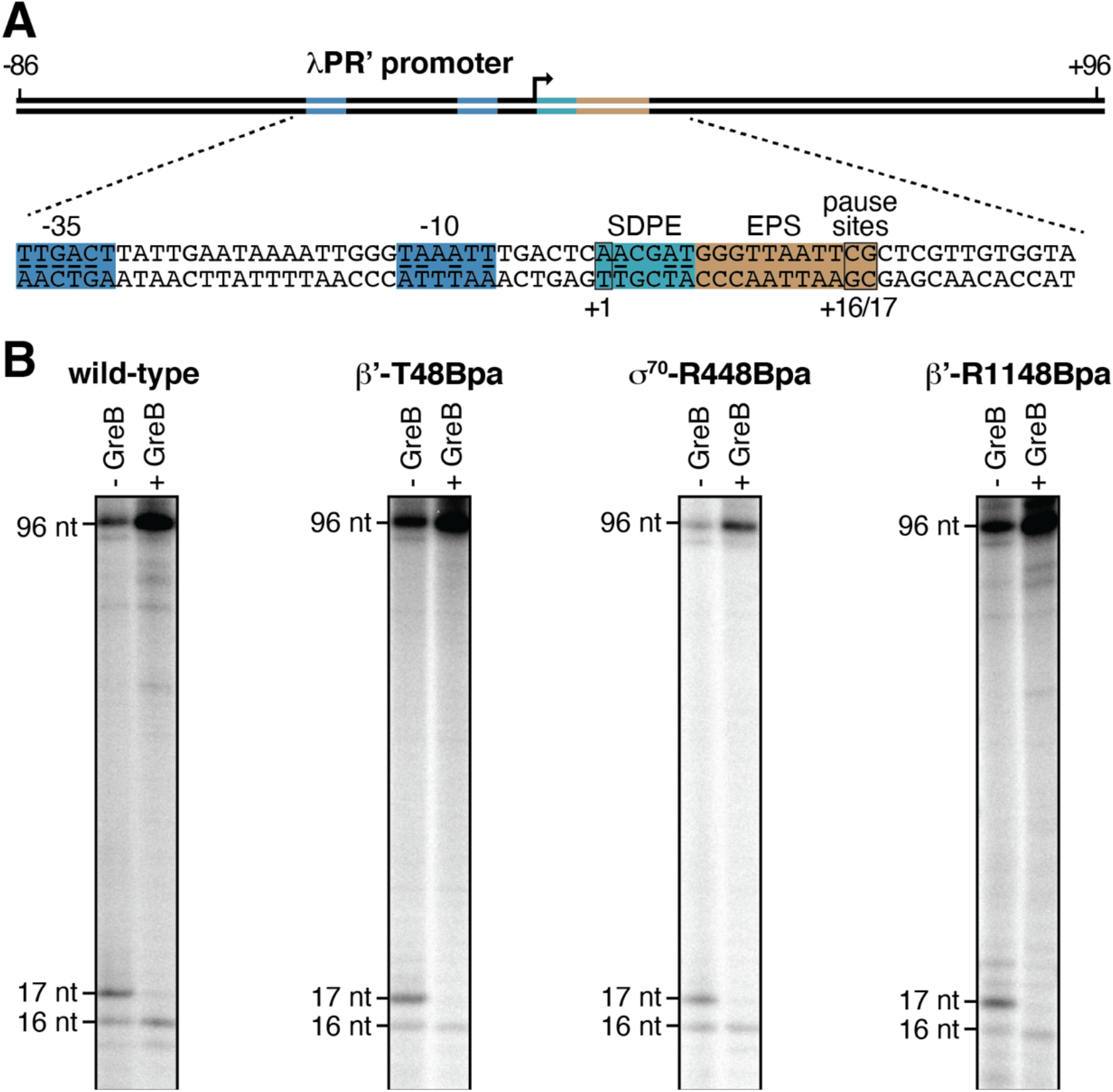
Use of site-specific protein-DNA photocrosslinking to define positions of RNAP trailing and leading edges and of σ relative to DNA at λPR’: pausing properties of Bpa-labeled RNAP and σ derivatives. (A) DNA template containing *λ*PR’ promoter. Colors as in Figure 1A. (B) Gel images of PAGE analysis of RNA products for *in vitro* transcription reactions performed with wild-type RNAP, RNAP-*β*’ ^T48Bpa^ (Bpa at RNAP trailing edge), RNAP- σ^70 R448Bpa^ (Bpa in σR2), and RNAP-*β*’ ^R1148Bpa^ (Bpa at RNAP leading edge). Positions of 16- and 17-nt RNA products and 96-nt full-length RNA products are indicated.

**Figure S2.**
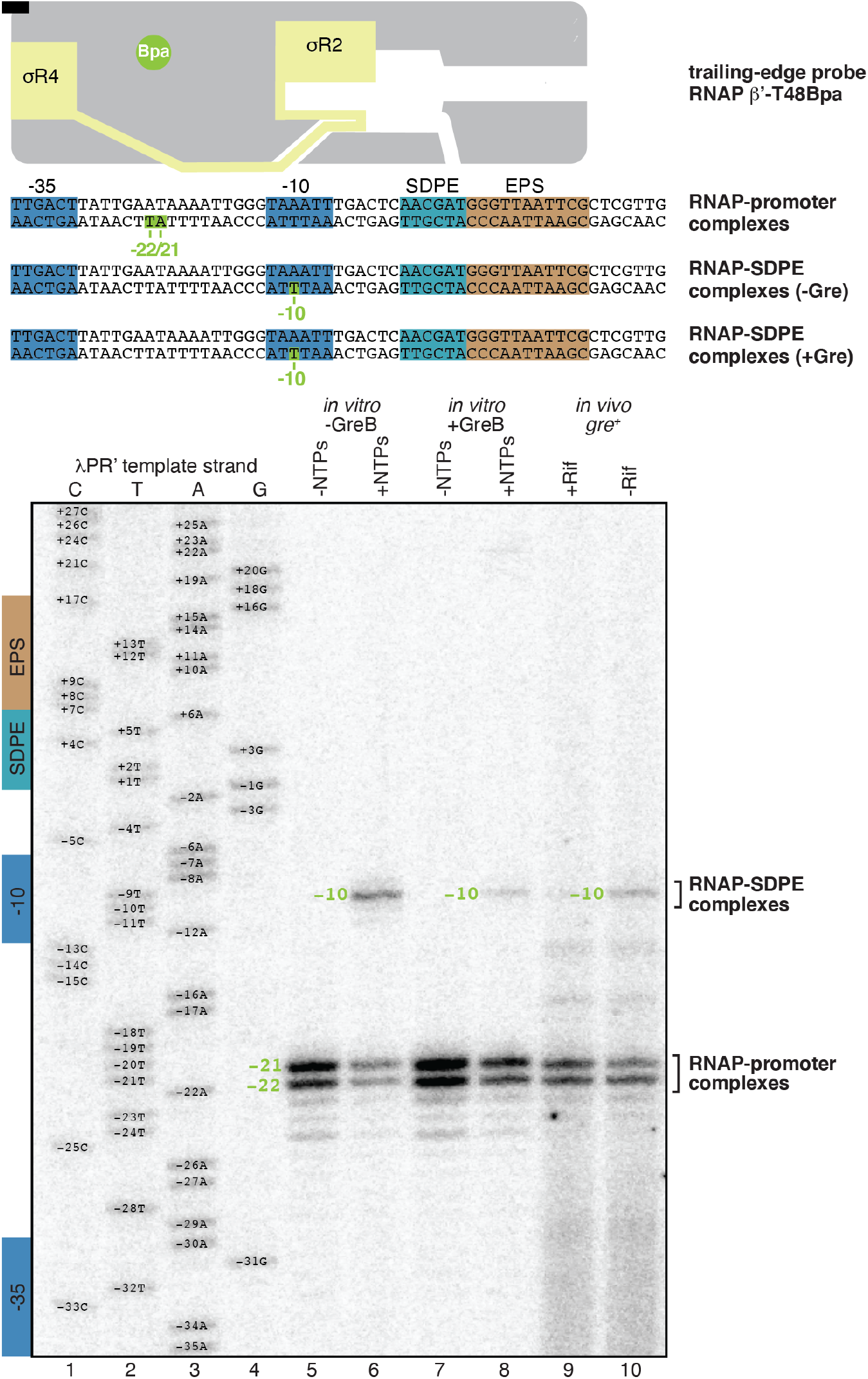
Use of site-specific protein-DNA photocrosslinking to define position of the RNAP trailing edge at λPR’. Top, *λ*PR’ promoter. Observed trailing-edge crosslinking sites are in olive green. Other colors as in Figure 1A. Bottom, positions of RNAP trailing edge in RNAP-promoter complexes or RNAP- SDPE complexes at *λ*PR’. Figure shows sequence ladder generated using *λ*PR’ (lanes 1-4) and primer-extension mapping of crosslinking sites for each experimental condition *in vitro* and *in vivo*, identified at top (lanes 5-10).

**Figure S3.**
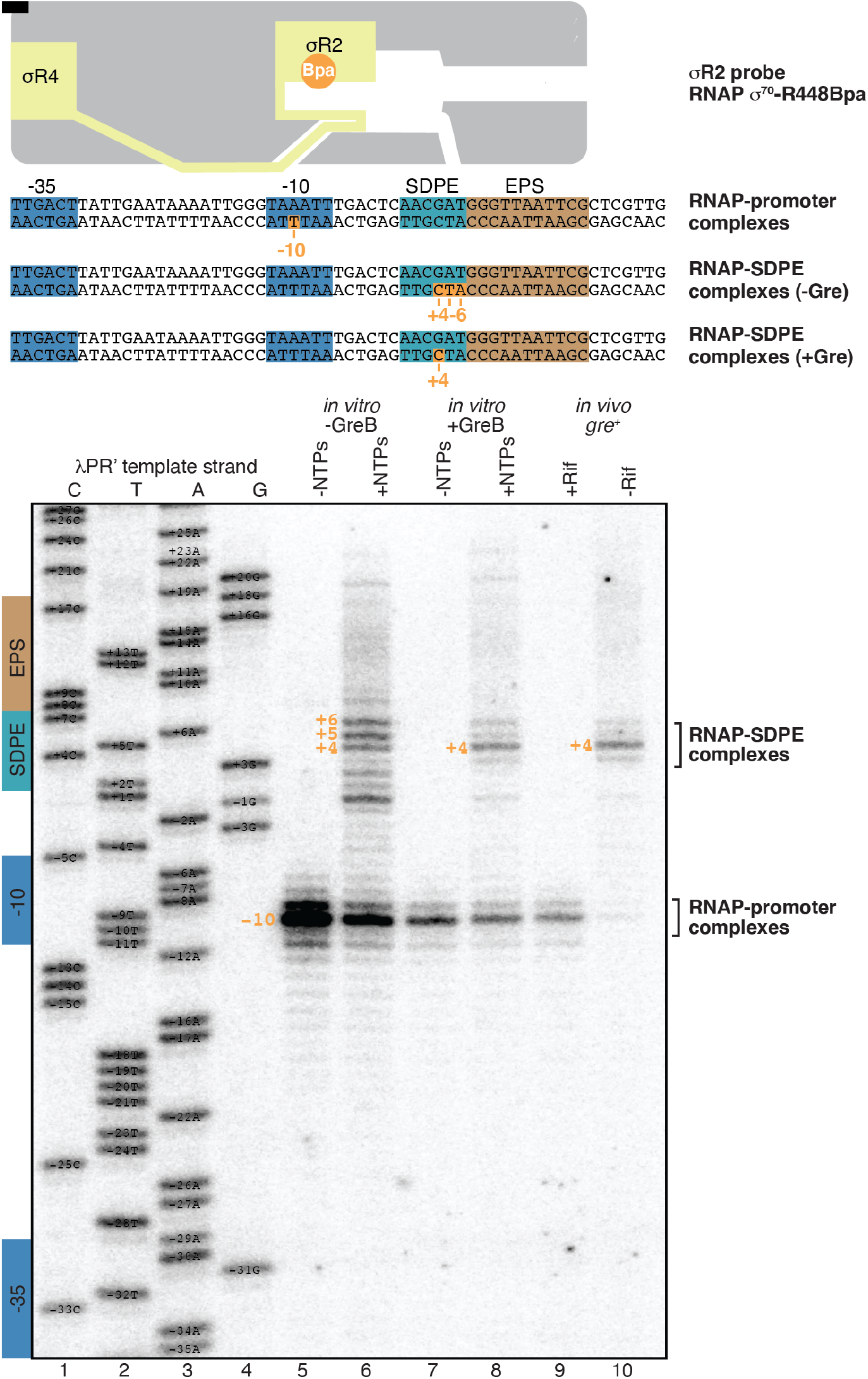
Use of site-specific protein-DNA photocrosslinking to define position of σR2 at λPR’: results. Top, *λ*PR’ promoter. Observed *σ*R2 crosslinking sites are in orange. Other colors as in Figure 1A. Bottom, positions of *σ*R2 in RNAP-promoter complexes or RNAP-SDPE complexes at *λ*PR’. Figure shows sequence ladder generated using *λ*PR’ (lanes 1-4) and primer-extension mapping of crosslinking sites for each experimental condition *in vitro* and *in vivo*, identified at top (lanes 5- 10).

**Figure S4.**
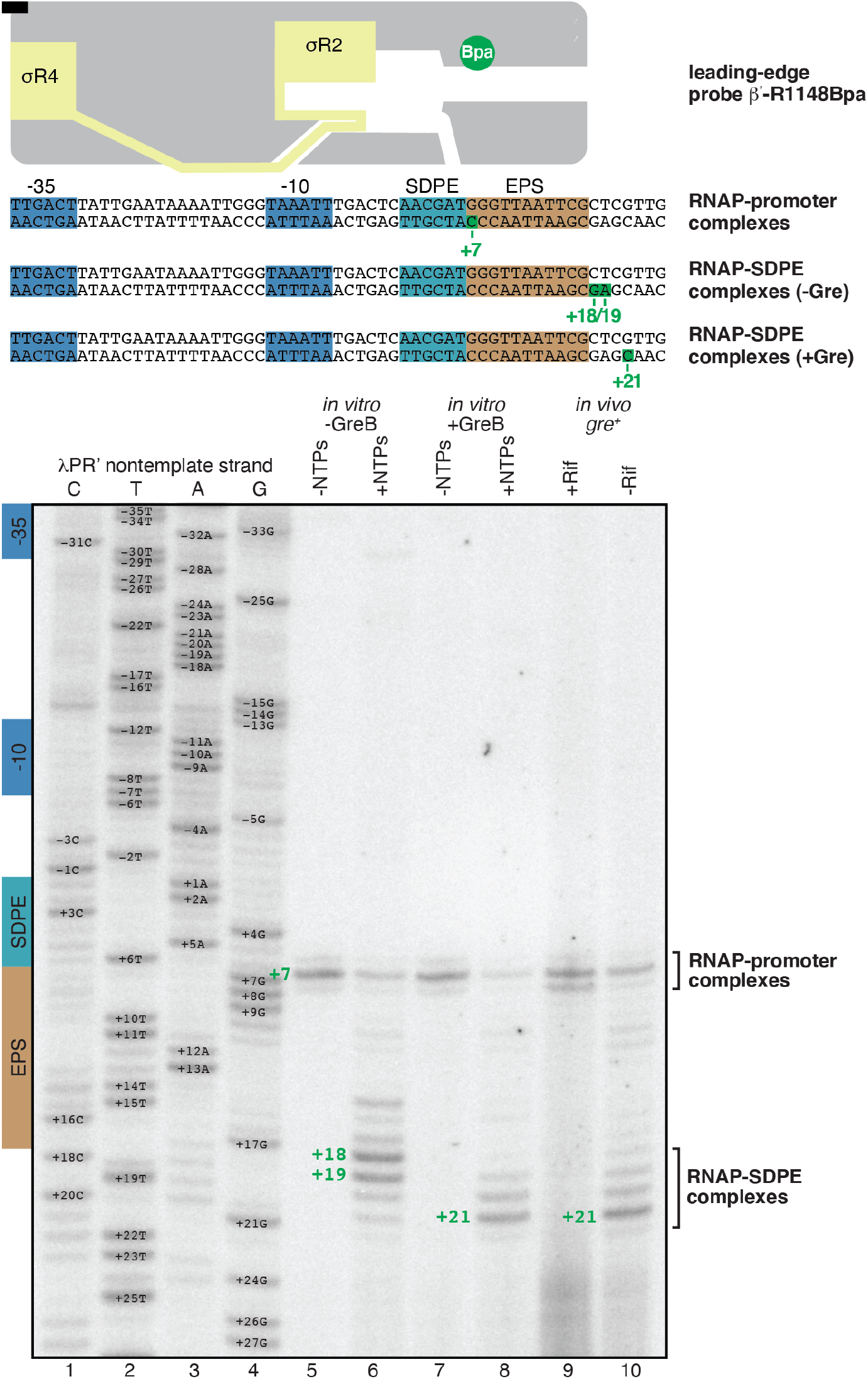
Use of site-specific protein-DNA photocrosslinking to define position of the RNAP leading edge at λPR’. Top, *λ*PR’ promoter. Observed leading-edge crosslinking sites are in forest green. Other colors as in Figure 1A. Bottom, positions of RNAP leading edge in RNAP-promoter complexes or RNAP- SDPE complexes at *λ*PR’. Figure shows sequence ladder generated using *λ*PR’ (lanes 1-4) and primer-extension mapping of crosslinking sites for each experimental condition *in vitro* and *in vivo*, identified at top (lanes 5-10).

**Figure S5.**
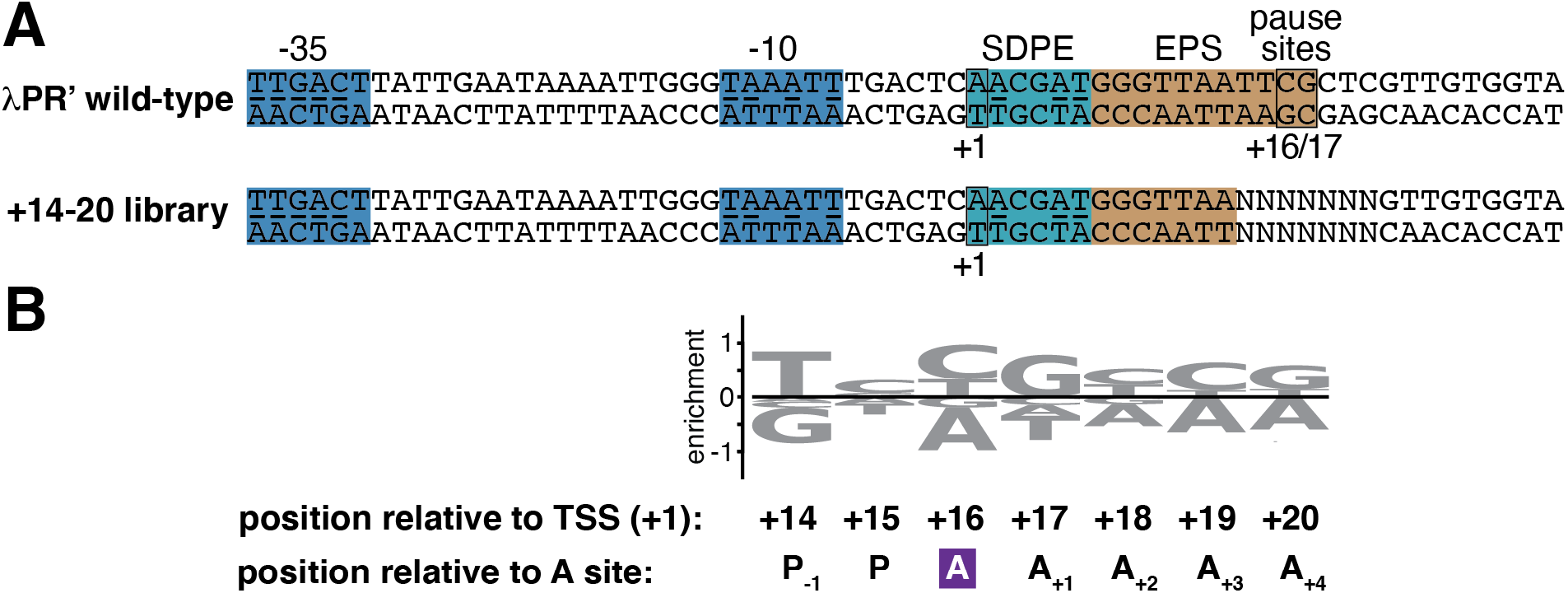
Sequence determinants for scrunching in σ-dependent pausing. (A) DNA templates containing wild-type *λ*PR’ or +14-20 library. NNNNNNN, randomized nucleotides of +14-20 library. Other colors as in Figure 1A. (B) Full sequence logo (positions +14 to +20) quantifying the formation and/or stability of major scrunched σ-dependent paused complex (RNAP-active-center A-site at position +16).

**Figure S6.**
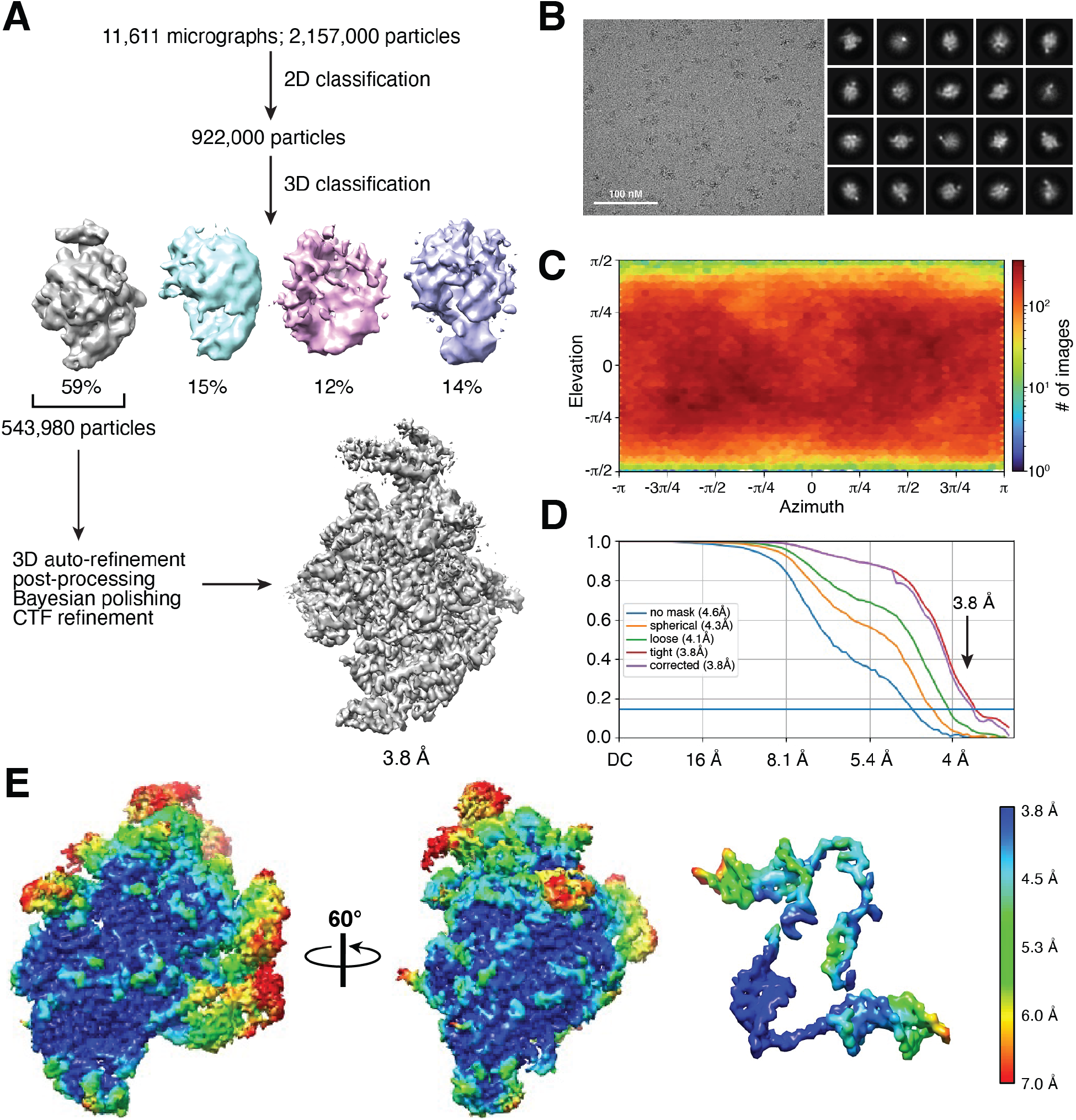
Structural basis for scrunching in σ-dependent pausing: cryo-EM structure determination. **(A)** Data processing scheme. **(B)** Representative electron micrograph (left; 100 nm scale bar) and representative class averages (right). **(C)** Orientational distribution. **(D)** Gold-standard Fourier shell correlation (GSFSC) resolution plot. **(E)** EM density maps colored by local resolution. Left, overall structure (two views; orientations as in Figure 6B). Right, DNA and RNA in structure (view orientation as in Figure 6B, left).

**Figure S7.**
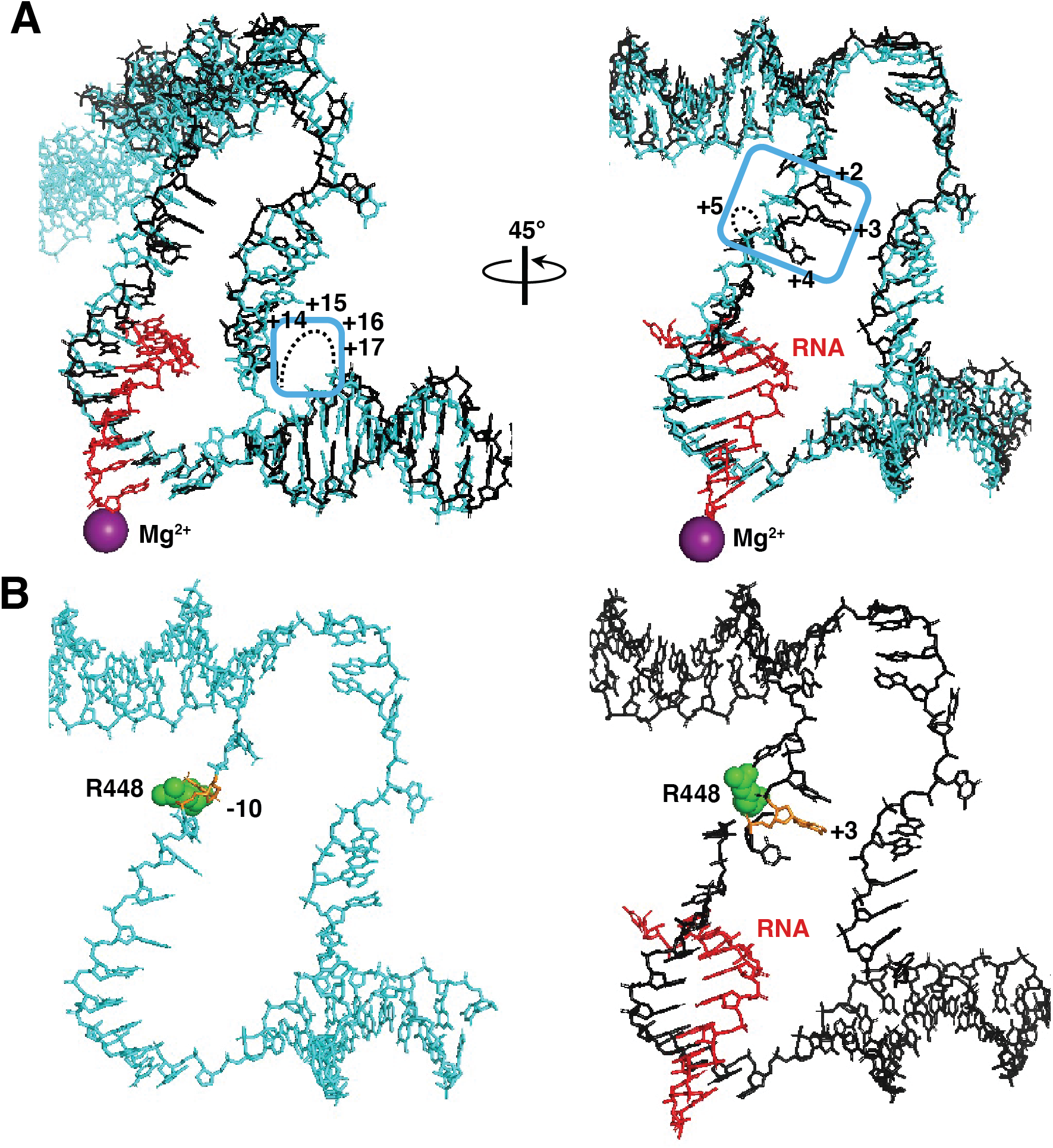
Structural basis for scrunching in σ-dependent pausing: scrunched nontemplate-strand and template-strand DNA nucleotides in pTEC. (A) Superimposition of DNA (black), RNA (red), and RNAP-active-center Mg^2+^ (violet sphere) in structure of pTEC (3 bp of scrunching; Figure 6) on DNA (cyan) in structure of RPo (no scrunching; (29); PDB 5I2D) nontemplate-strand (left) and template-strand (right) DNA nucleotides in pTEC (view orientations as in Figure 6C). Blue boxes, DNA nucleotides disordered or repositioned due to DNA scrunching; black dots, DNA nucleotides disordered due to DNA scrunching. Nucleotides numbered as in *λ*PR’. (B) Comparison of relative positions of *σ*R2 R448 and template strand of -10 element in RPo (left; (29); PDB 5I2D) to relative positions of *σ*R2 R448 and template strand of SDPE in pTEC (right; Figure 6). Green, *σ*R2 R448; orange, template-strand nucleotide at third position of -10 element (left, *λ*PR’ position -10) or template-strand nucleotide at third position of SDPE (right, *λ*PR’ position +3). View orientation and other colors as in panel A, left.

**Table S1.**
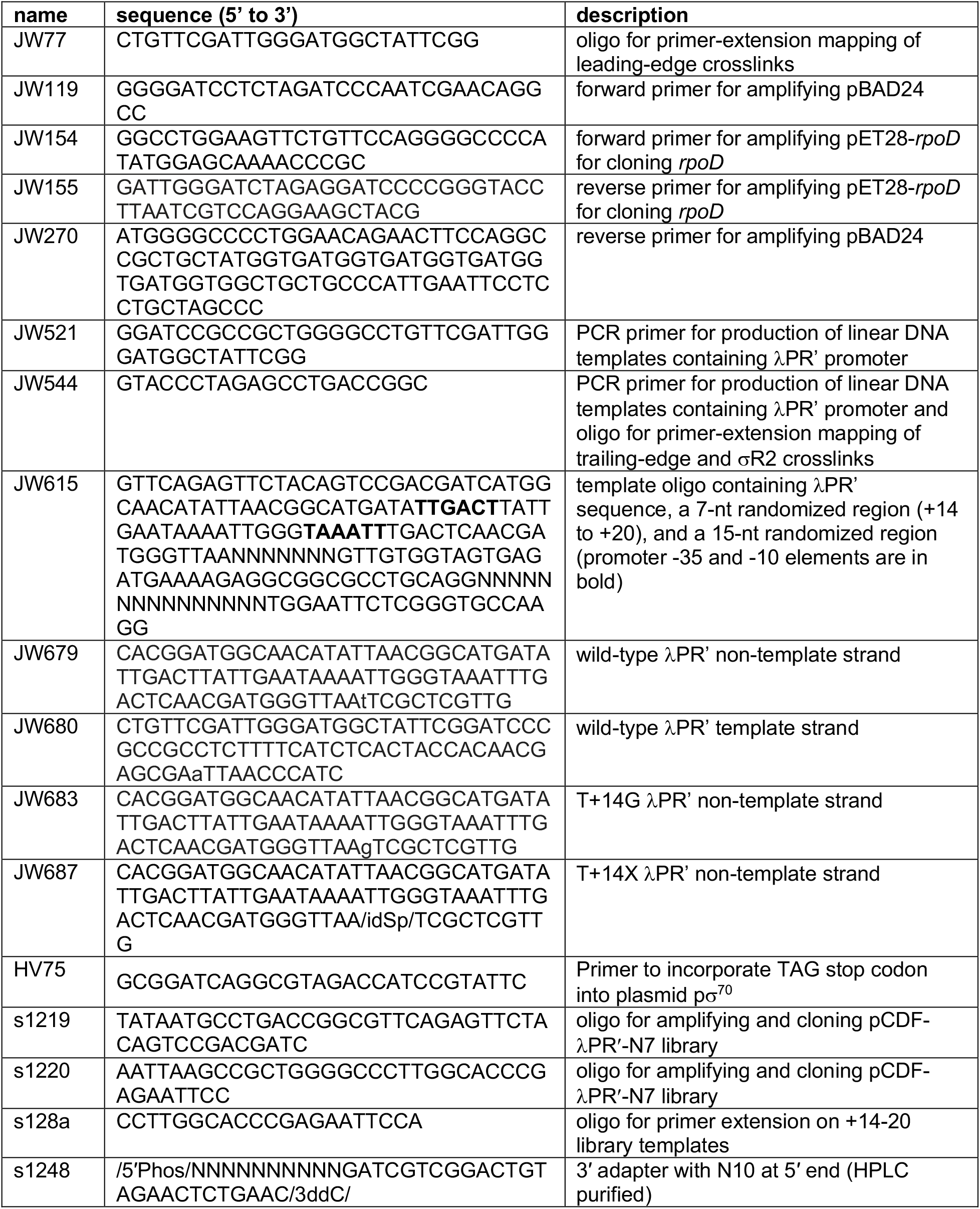
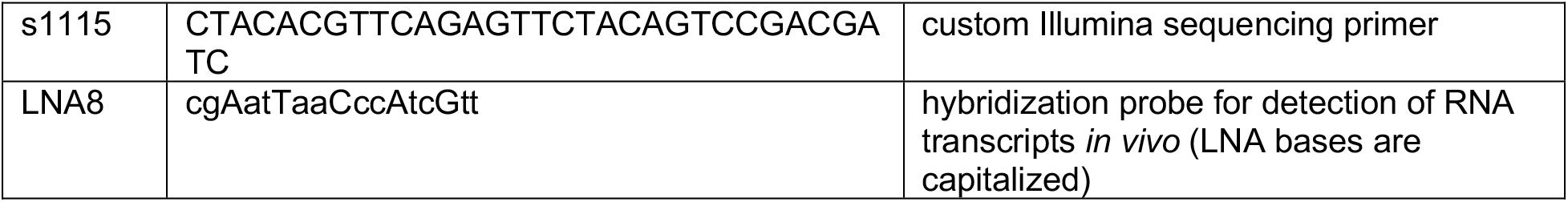
Oligonucleotides.

